# Disease mutations in the PWWP domain of DNMT3A affect chromatin recruitment through multiple mechanisms

**DOI:** 10.64898/2026.08.28.747843

**Authors:** Hannah Wapenaar, Gillian Clifford, Francesca Taglini, Finlay McGhie, Willow Rolls, Yujie Zhang, Duncan Sproul, Marcus D. Wilson

## Abstract

DNMT3A is a *de novo* DNA methyltransferase whose recruitment to chromatin regulates its function. Missense mutations within the chromatin-binding PWWP domain are associated with diverse human disorders, yet how mutations in the same domain produce distinct phenotypes remains unclear. Here we systematically characterise 19 clinically reported mutations in the PWWP domain of DNMT3A that are associated with Heyn-Sproul-Jackson syndrome (HESJAS), paraganglioma (PG) and clonal haematopoiesis (CH). We show that all PWWP-domain mutations associated with HESJAS abolished interaction with H3K36me2 modified nucleosomes, defining this as a consistent biochemical feature of HESJAS. In contrast, mutations from all disease classes differentially altered DNA binding of the PWWP domain, driven by alterations in the net charge of the domain. However, these effects are largely overcome by inclusion of the DNNMT3A1 N-terminal region, which is absent from its embryonic isoform, suggesting that PWWP mutations may differentially affect DNMT3A function through development. Changes in the thermal stability of the isolated PWWP domain mutants did not directly translate into altered stability of full-length DNMT3A1 in cells. We show that HESJAS mutations can affect the intramolecular interaction between the PWWP and adjacent ADD domain, an interaction proposed to contribute to the autoinhibitory function of the ADD domain. However, not all mutations behaved in the same way, suggesting that multiple factors govern the intramolecular autoinhibition of DNMT3A. Together, this study advances our understanding of the molecular mechanisms by which DNMT3A PWWP-domain mutations are mechanistically heterogeneous, providing a biochemical framework that contributes to distinct disease phenotypes.

**Highlights:**

- Mutations in the DNMT3A PWWP domain are associated with disease phenotypes
- Loss of H3K36me2 binding is a unifying mechanism for HESJAS
- The N-terminal extension of DNMT3A1 alters isoform DNA binding
- PWWP mutations modify the interaction with the ADD domain of DNMT3A

## Introduction

DNMT3A1 is a *de novo* DNA methyltransferase important for establishing correct DNA methylation patterns in cells [1]. Accurate recruitment and regulation of DNMT3A to chromatin is essential for normal epigenetic processes. DNMT3A consists of a catalytic C-terminal domain and a regulatory N-terminal section that integrates multiple chromatin-binding activities. These include a catalytic-autoinhibitory ATRX-DNMT3-DNMT3L (ADD) domain that recognises unmodified histone H3 at lysine 4 (H3K4) [2, 3], and a Pro-Trp-Trp-Pro (PWWP) domain that binds methylated H3K36 [4–7]. DNMT3A is expressed as two developmentally regulated isoforms generated through alternative promoter usage (Figure 1A) [8, 9]. The longer isoform, DNMT3A1, predominates in somatic cells and contains an extended, largely disordered N-terminal region that harbours a ubiquitin-dependent recruitment (UDR) motif that contacts the nucleosome surface and histone ubiquitylation [10–13]. The N-terminal region also contains DNA-binding activity [14, 15]. The shorter isoform, DNMT3A2, is highly expressed during embryonic development and lacks this N-terminal region (Figure 1A) [8–10, 16, 17], making chromatin recruitment mechanisms involving this region specific to DNMT3A1.

**Figure 1.**
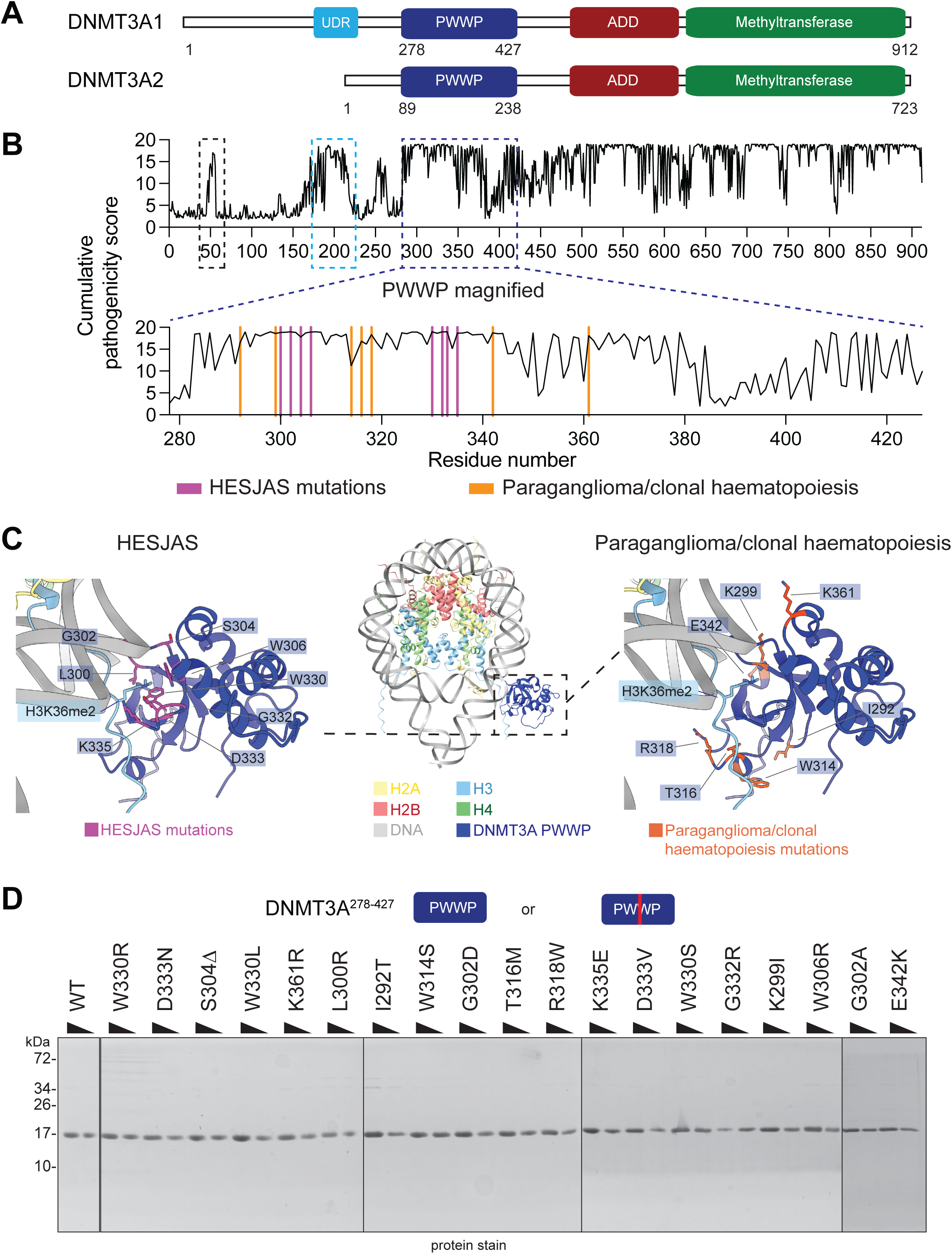
Overview of DNMT3A and the PWWP-domain disease mutations analysed in this study. A. Schematic overview of DNMT3A1 containing a methyltransferase domain (green), ATRX-DNMT3-DNMT3L (ADD, red) a Pro-Trp-Trp-Pro (PWWP, blue) domain N-terminal region containing a ubiquitin-dependent recruitment motif (UDR, cyan) and isoform DNMT3A2, which lacks the N-terminal region and UDR. B. Cumulative predicted pathogenicity scores from AlphaMissense for full-length DNMT3A1 (top, region around 50 AA in black box, UDR in light blue) and the DNMT3A1 PWWP domain (residues 278-427, dark blue box, bottom). Heyn-Sproul-Jackson syndrome (HESJAS) mutations (magenta) and Paraganglioma/clonal haematopoiesis mutations (orange) analysed in this study are indicated. C. AlphaFold 3 model of DNMT3A1 PWWP domain (blue) bound to an H3K36me2 nucleosome wrapped with 175 bp Widom 601 DNA (middle). Left: HESJAS mutations used in this study (magenta) mapped on the model, right: paraganglioma and clonal haematopoiesis used in this study (orange) mapped on the model. D. SDS-PAGE gel of 1 and 0.5 μg of purified DNMT3A1^278-427^ wild type (WT) and mutant proteins, stained with colloidal Coomassie stain.

Heterozygous mutations in DNMT3A are implicated in a range of human disorders. These include clonal haematopoiesis (CH) [18–20], acute myeloid leukaemia (AML) [21–24] and paraganglioma (PG) [24, 25]. Single missense mutations are also drivers in developmental disorders including Tatton-Brown-Rahman syndrome (TBRS) [26] and Heyn-Sproul-Jackson syndrome (HESJAS) [27, 28]. While many disease-associated mutations reduce protein stability or catalytic activity - particularly in TBRS [29–31] and AML [29–31] - a distinct subset of mutations preserve protein stability and do not directly affect the catalytic domain of DNMT3A [32, 33]. Strikingly, many of the mutations conserving stability and activity, cluster within the PWWP domain of DNMT3A and are associated with distinct disorders, namely HESJAS [27, 28, 34–36], paraganglioma [25] and clonal haematopoiesis [32, 37]. HESJAS is a developmental disorder characterised by microcephalic dwarfism and intellectual disability, and has recently been reclassified as a progeroid syndrome [27, 28]. In contrast, paraganglioma-associated mutations predispose to tumour formation, although several PWWP-mutations produce gain-of-function phenotypes resembling HESJAS [25] and there is emerging clinical overlap between the two disorders, including reports of paragangliomas in HESJAS patients [27, 28]. Even within HESJAS, individual PWWP mutations produce a spectrum of clinical severity [28]. Together, these observations suggest that disease-associated PWWP mutations are likely to perturb multiple biochemical functions. However, which mechanisms are responsible for causing such a variety of phenotypes from a single domain is incompletely understood.

Previous work has introduced clinical mutations to inform on DNMT3A function in cell-biological and biochemical studies; a two-way street that has also ultimately informed on disease pathologies [33]. The HESJAS W330R mutation disrupts H3K36me2 interaction and leads to redistribution of DNMT3A1 to H2AK119ub-enriched regions of the genome, driven via the UDR in DNMT3A1’s N-terminal region [10–13, 35]. This redistribution produces DNA methylation changes and HESJAS-like phenotypes in mouse models [27, 28, 38]. However, both targeted PWWP mutational analyses analysis [32, 35] and unbiased base editor scanning [37] have demonstrated mutation-specific differential effects on DNA methylation in cells.

In addition to recognising H3K36 methylation, the PWWP domain binds DNA [7, 37, 39]. Both HESJAS and paraganglioma mutations in this domain have shown to affect DNA binding [37, 40]. Furthermore, the PWWP domain has recently been proposed to participate in intramolecular autoinhibition by contacting the ADD domain, with certain mutations perturbing this interaction [40]. An analogous regulatory mechanism has been described for the paralogue DNMT3B [41, 42]. Together, these findings suggest that distinct disease-associated mutations may perturb different biochemical functions of the PWWP domain, leading to different disease phenotypes.

To determine whether clinically distinct PWWP mutations have distinct biochemical consequences, we systematically characterised 19 clinically reported PWWP-domain mutations linked to HESJAS, paraganglioma or clonal haematopoiesis (Table 1). We found that the mutations displayed diverse effects on the biological roles of the isolated domain, but some effects were ameliorated in larger constructs. However, we find every HESJAS-associated mutation reduced binding to recombinant H3K36me2 modified nucleosome core particles, which we propose biochemically underlies the central pathogenesis of this disorder.

**Table 1.**
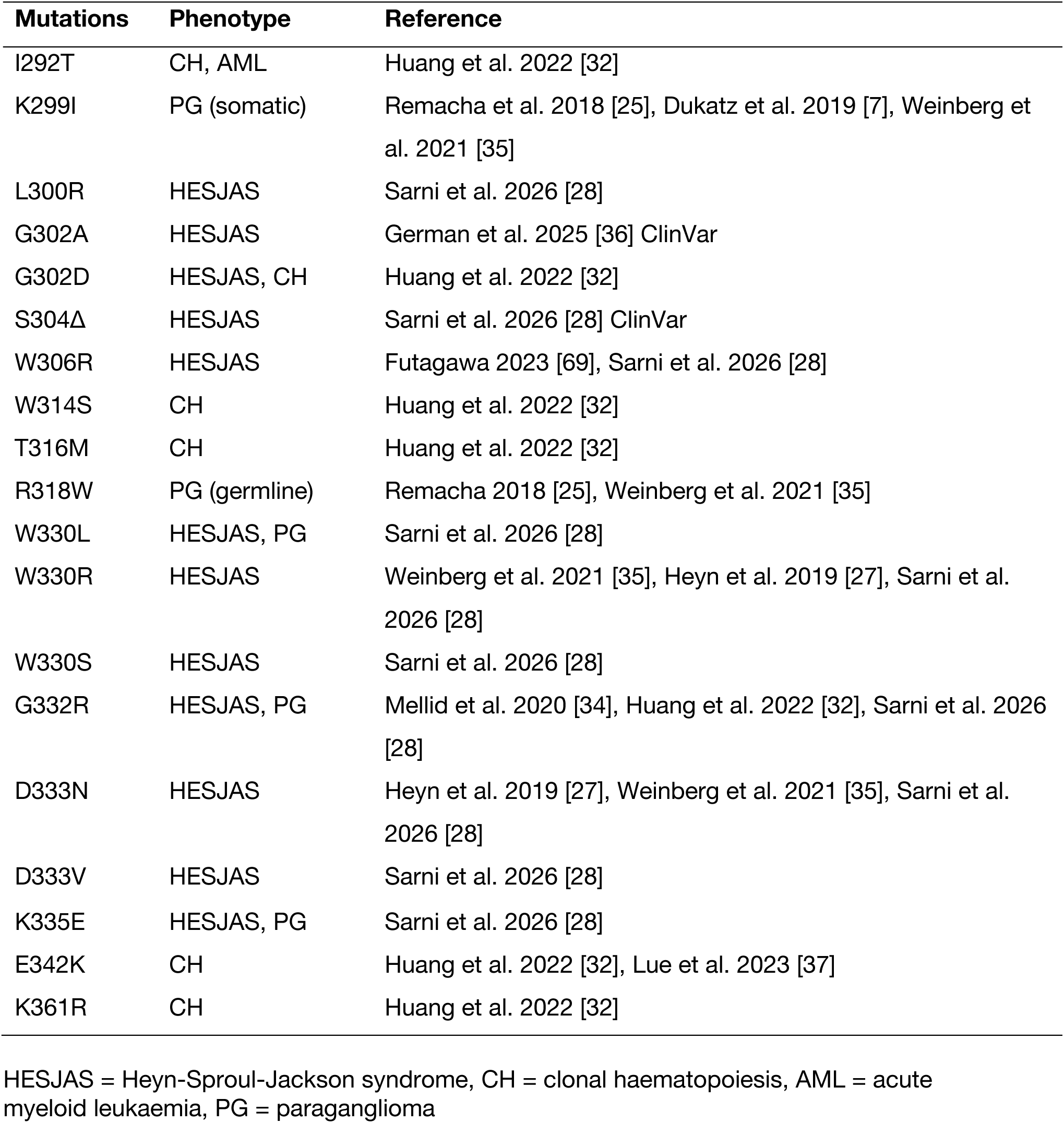
Overview of disease mutations in the PWWP domain of DNMT3A1 used in this study.

## Results

### Mutations in the PWWP domain are spatially distinct in different disorders

A number of DNMT3A PWWP-domain mutations have been linked to human disease [25, 27, 28, 32, 33]. To explore the biochemical link between DNMT3A PWWP mutations, function and whether this could be corelated with disease states, we selected 19 clinically reported patient mutations associated with HESJAS, paraganglioma or clonal haematopoiesis for systematic analysis (Table 1). We excluded mutations affecting TBRS and AML which commonly affect protein stability or co-occur with other mutations. Selected mutations were either single base missense changes leading to residue changes or in the case of S304Δ loss of three bases leading to one fewer amino acid. Some residues are hotspots for mutation with 2 or 3 variant amino acids identified in patients (G302, D333 and W330 respectively).

To place these mutations in the context of the full DNMT3A1 protein, we bioinformatically explored predicted pathogenicity [43] across all possible substitutions at each residue of DNMT3A1 (Figure 1B). The disordered N-terminal region was largely predicted to tolerate mutation, with only a few high-scoring regions, including the previous described UDR motif and an additional peak around residue 50 (Figure 1B, top, black box). By contrast, the PWWP, ADD and catalytic domains were predicted to be highly sensitive to mutation, and all of the selected diseases-associated PWWP-domain residues fell within regions of high predicted pathogenicity (Figure 1B, bottom PWWP magnifcation).

To visually map disease mutations onto the PWWP domains chromatin-binding surface, we generated a prediction of the DNMT3A PWWP domain (aa 278-427, dark blue) bound to an H3K36me2 modified nucleosome using AlphaFold 3 [44] (Figure 1C, middle). All generated models adopted a similar binding mode with comparable confidence scores, differing mainly in their flexible DNA linker arms and histone tails (Supp. Figure 1A&B). The PWWP domain was predicted to engage the nucleosome in a manner resembling the H3K36me3-reading PWWP domain of LEDGF [45, 46] with the H3K36me2 mark accommodated in the binding pocket analogous to that seen in the structure of the DNMT3B PWWP domain complexed to a H3K36me3 peptide [5]. All HESJAS missense mutations (Figure 1C left, magenta) mapped to the predicted binding site of H3K36me2, either within or adjacent to the methyl-lysine binding aromatic cage [28]. In contrast, paraganglioma and clonal haematopoiesis point mutations were located distal to this binding site (Figure 1C right, orange), but still clustered on the nucleosome adjacent face of the PWWP domain, suggesting potential involvement in nucleosome interaction. We therefore hypothesised that HESJAS substitution mutations, but not paraganglioma or clonal haematopoiesis mutations, would specifically impair H3K36me2 recognition in the context of a nucleosome.

### HESJAS mutations impair binding to H3K36me2 modified nucleosomes

To test this hypothesis, we recombinantly expressed and purified the PWWP domain of DNMT3A (DNMT3A1^278-427^). Wild type (WT) and 19 identified missense mutants were purified similarly (Figure 1D) and molecular weight verified by 1D intact-protein mass spectrometry (Supp. Table 1). We wrapped defined nucleosome core particles (NCPs) with 145bp of fluorescently labelled strong nucleosome positioning Widom 601 DNA [47]. Using fluorescence polarisation, we first compared WT PWWP domain binding to NCPs that were either unmodified or contained a chemically installed H3K36me2 modification (Supp. Figure 1C). The WT PWWP domain showed strong preference for H3K36me2 modified NCPs over unmodified NCPs under these conditions (Figure 2A), suggesting this was a suitable assay to screen for H3K36me2 specificity of the purified mutants.

**Figure 2.**
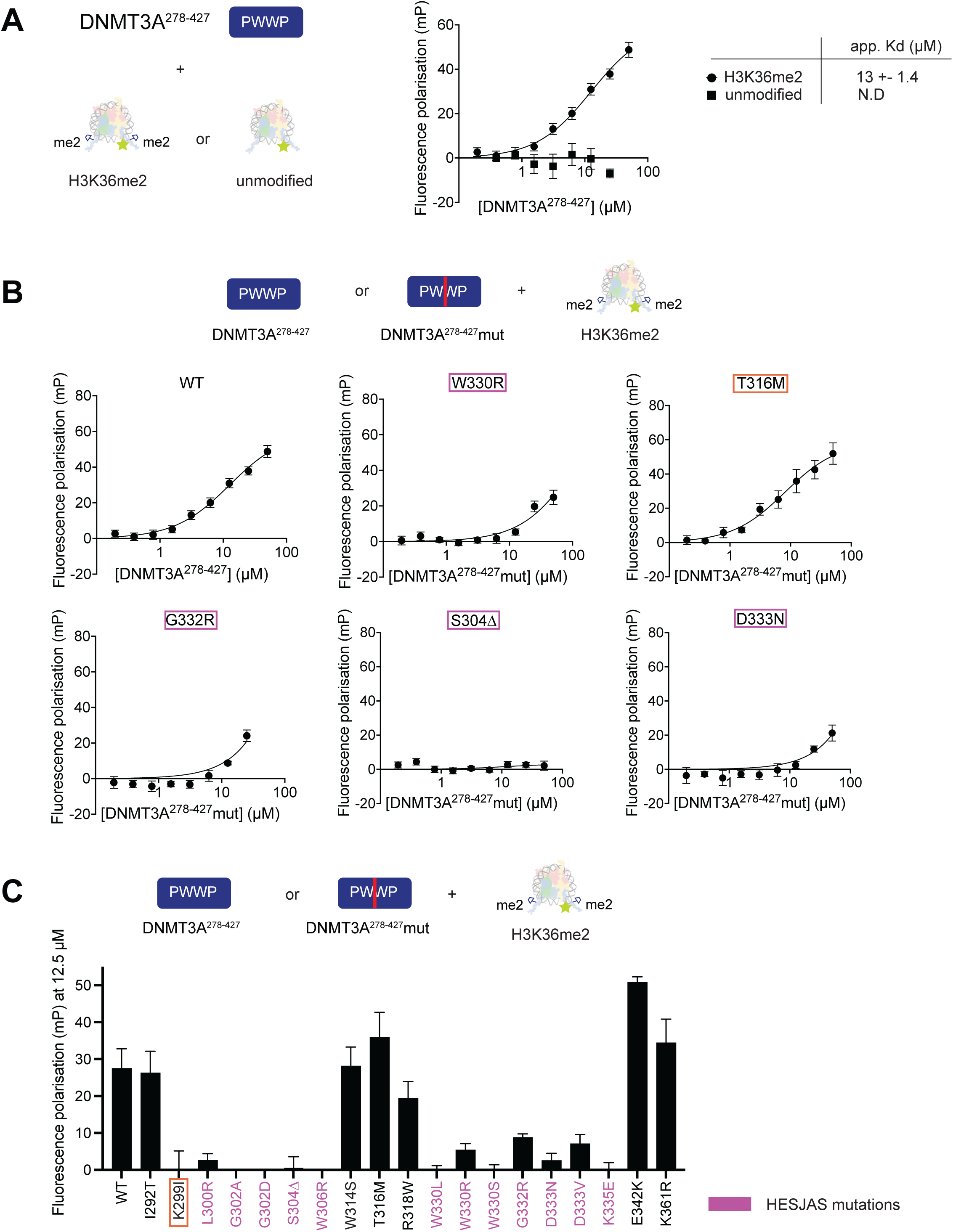
HESJAS mutations inhibit binding to the H3K36me2 mark on nucleosomes. A. Fluorescent polarisation assay investigating binding of DNMT3A1^278-427^ to unmodified or H3K36me2 modified nucleosomes. DNMT3A1^278-427^ (0-25 μM, 2x dilution series) was incubated with nucleosomes wrapped with 5′ 6-FAM- labelled 145bp Widom601 DNA (6.7 nM). Apparent K_D_ values are indicated (N.D., not determined), data derived from three independent experiments. B. Representative binding curves for selected DNMT3A1^278-427^ mutants (HESJAS mutations in magenta, clonal haematopoiesis in orange) to H3K36me2 modified nucleosomes by fluorescence polarisation assay. Curves for all mutants are shown in Supp. Figure 2; apparent K_D_ values are listed in Table 2. C. Summary of fluorescence polarisation values of DNMT3A1^278-427^ variants binding to H3K36me2 modified nucleosomes at 12.5 μM PWWP domain. Data taken from B and Supp. Figure 2. HESJAS mutations shown in magenta, paraganglioma mutation K299I marked with orange box.

**Table 2.**
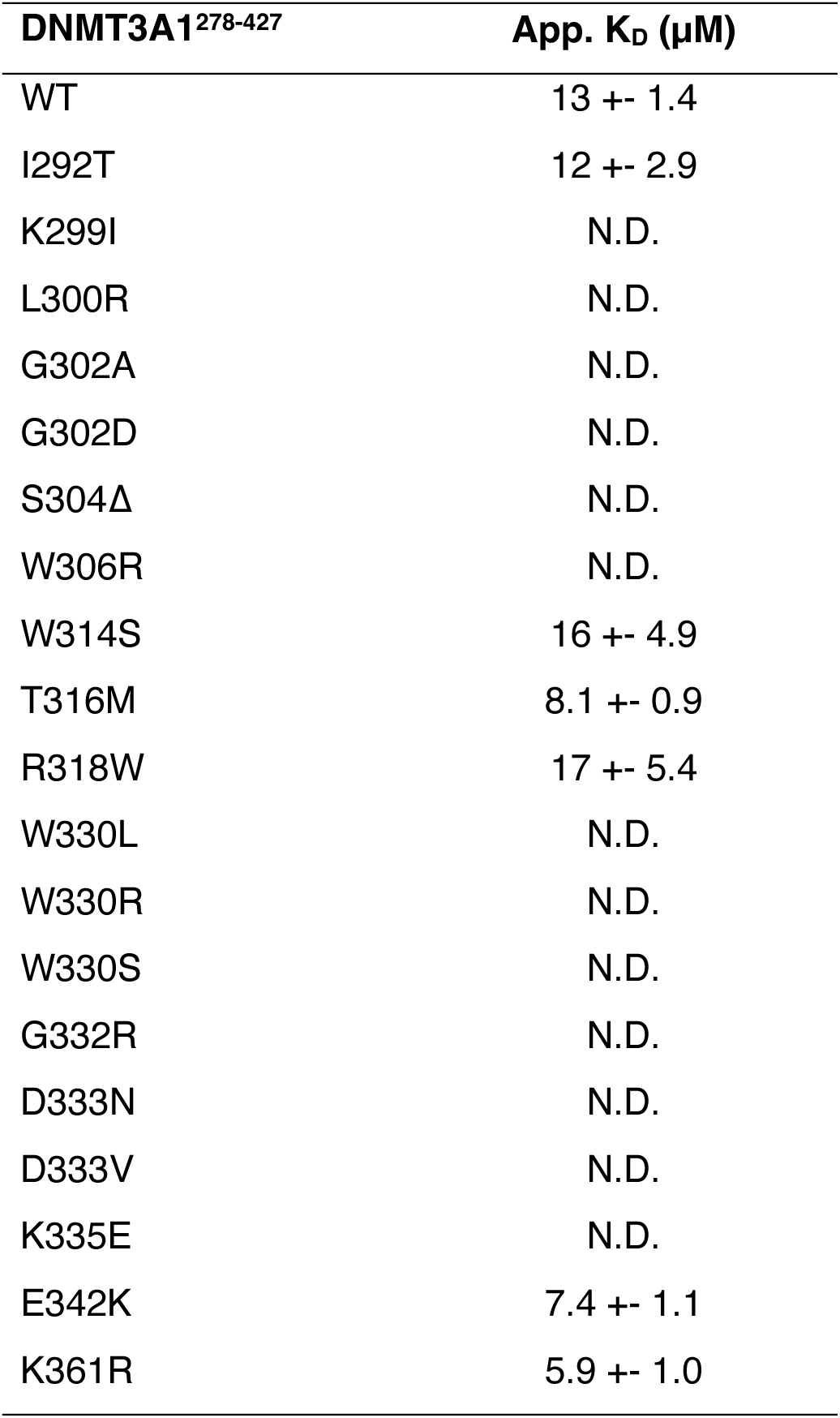
Apparent K_D_ values of DNMT3A1^278-427^ binding to H3K36me2 modified nucleosomes determined by fluorescence polarisation.

We then systematically screened all mutants for binding to H3K36me2 modified NCPs (Figure 2B&C, Supp. Figure 2, Table 2 of derived dissociation constants, K_D_ values). All HESJAS-associated mutations showed a strong reduction in affinity for H3K36me2 modified NCPs (Figure 2C, magenta), leading polarisation changes that were below the levels of quantifiable curve fitting, even at high PWWP concentrations. This was true even for subtle chemical alterations such as G302A mutation. However, not all mutations were equally poor binders; for example, substitution of tryptophan at position 330 for leucine or serine showed no measurable binding but a detectable dose response of polarisation was observed for an arginine substitution (Supp. Figure 2). This suggests that the exact chemical nature of residues rather than just their mutation affects interaction. In contrast, mutations solely associated with paraganglioma or clonal haematopoiesis largely retained WT-like binding, albeit with slightly altered fit K_D_ values (Table 2). This suggests that despite similar location on the PWWP domain H3K36me2-binding is not a molecular defect for these conditions. Paraganglioma mutation K299I was the one exception to this trend, showing diminished H3K36me2 binding, consistent with previous reports [7]. Notably, K299I arose as a somatic mutation in all described paraganglioma patients [25], which may explain the absence of a reported developmental HESJAS phenotype in these individuals. Together, these results indicate that loss of H3K36me2 binding is a shared feature of HESJAS-associated mutations, and suggest that germline mutations that ablate H3K36me2 recognition would give rise to HESJAS. Because paraganglioma- and clonal haematopoiesis-associated mutations do not uniformly affect H3K36me2 binding, additional mechanisms are likely to contribute to these conditions.

### The effect of PWWP mutations on DNA binding is buffered by the DNMT3A1 N-terminal region

Despite reduced binding affinity, some HESJAS mutations still showed some observable response in polarisation assays (e.g. W330R and G332R in Figure 2B). Furthermore, the E342K mutation showed a slight but consistent increase in overall binding affinity to nucleosomes, suggesting a gain-of-function perhaps mediated by recognition of nucleosomal features other than H3K36me2. To test general nucleosome binding we used electrophoretic mobility shift assays (EMSAs) as an orthologous assay to fluorescence polarisation. The WT PWWP domain indeed bound unmodified nucleosomes, with a preference for H3K36me2 modified nucleosomes (Figure 3A&B), as reported previously [13]. Consistent with the fluorescence polarisation data, two HESJAS mutations S304Δ and D333N lost specificity for H3K36me2 modified nucleosomes (Figure 3A&B). Unexpectedly, however, both mutations also altered the affinity to unmodified nucleosomes: D333N increased binding relative to WT, whereas S304Δ decreased it (Figure 3A&B).

**Figure 3.**
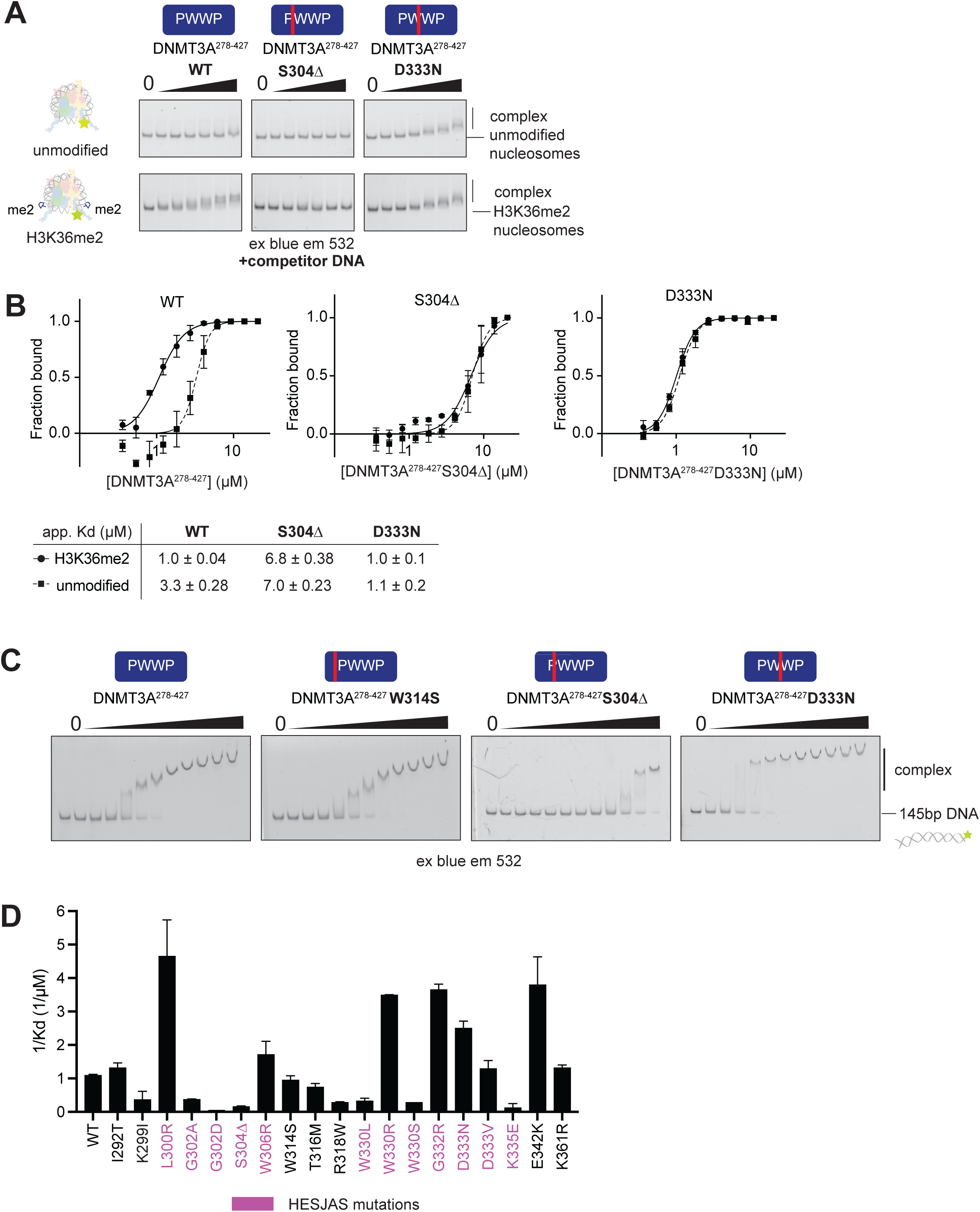
PWWP-domain mutations modulate DNA binding in a charge-dependent manner. A. EMSAs comparing the binding of WT, S304Δ and D333N DNMT3A1^278-427^ to unmodified and H3K36me2 modified nucleosome core particles (NCPs; 2.7 nM; 5′-6- FAM-labelled 145 bp Widom 601 DNA). PWWP protein (0-25 μM, 1.5-fold series); 0–3.3 µM shown for clarity. Complexes were resolved on native PAGE gels and imaged using blue light, emission 532 nm. Two replicates. B. Quantification of (A) from the full dilution series C. Representative EMSAs of DNMT3A1^278-427^ WT and mutants binding to 5′ 6-FAM-labelled 145bp Widom601 DNA. All mutants shown in Supp. Figure 3 and quantifications shown in Supp. Figure 4A. Experiments were repeated twice except for W330S done only once due to limited amount of material. D. Inverted K_D_ values (1/K_D_; higher value = tighter binding) of DNMT3A1^278-427^ WT and mutants binding to 5′ 6-FAM- labelled 145bp Widom601 DNA from C and Supp. Figure 4A. HESJAS mutations shown in magenta.

EMSA assays are generally sensitive to electrostatic interactions with DNA due to the lack of charge-neutralising salt in the resolving native gels, which may contribute to non-H3K36me2 nucleosome interaction. Indeed, the PWWP domains of DNMT3A and DNMT3B have been reported to bind to DNA [7, 39, 48, 49]. Therefore, we next tested binding of the PWWP domain mutants to a fluorescently labelled 145bp DNA duplex (Figure 3C&D, Supp Figure 3 & Supp Figure 4A). The majority of mutations affected DNA binding, unsurprising given the mutations cluster to the predicted nucleosomal-DNA contacting face of the PWWP (Figure 1C). However, DNA binding affinity varied widely between mutants, ranging from more than 4-fold tighter binding to a reduction below measurable in our assays. However, this pattern did not follow a clear correlative pattern with particular disease type or phenotype (Figure 3D, Table 3). Instead, DNA interaction tracked closely with the change in net charge imparted by each mutation: substitutions that added positive charge or removed negative charge (e.g. D333N, W330R, E342K, G332R) increased DNA binding, whereas others reduced interaction. Loss of affinity was observed for charge neutral mutations, likely due allosteric disruption of DNA interaction interfaces. This charge dependence is consistent with previous reports for W330R and E342K [37], K299I [7] and a related mutation associated with AML, K299N [37]. This suggests that PWWP-domain mutations can modulate DNA binding, but this change of electrostatic surface is not a diagnostic biochemical hallmark of specific DNMT3A-linked disease.

**Table 3.**
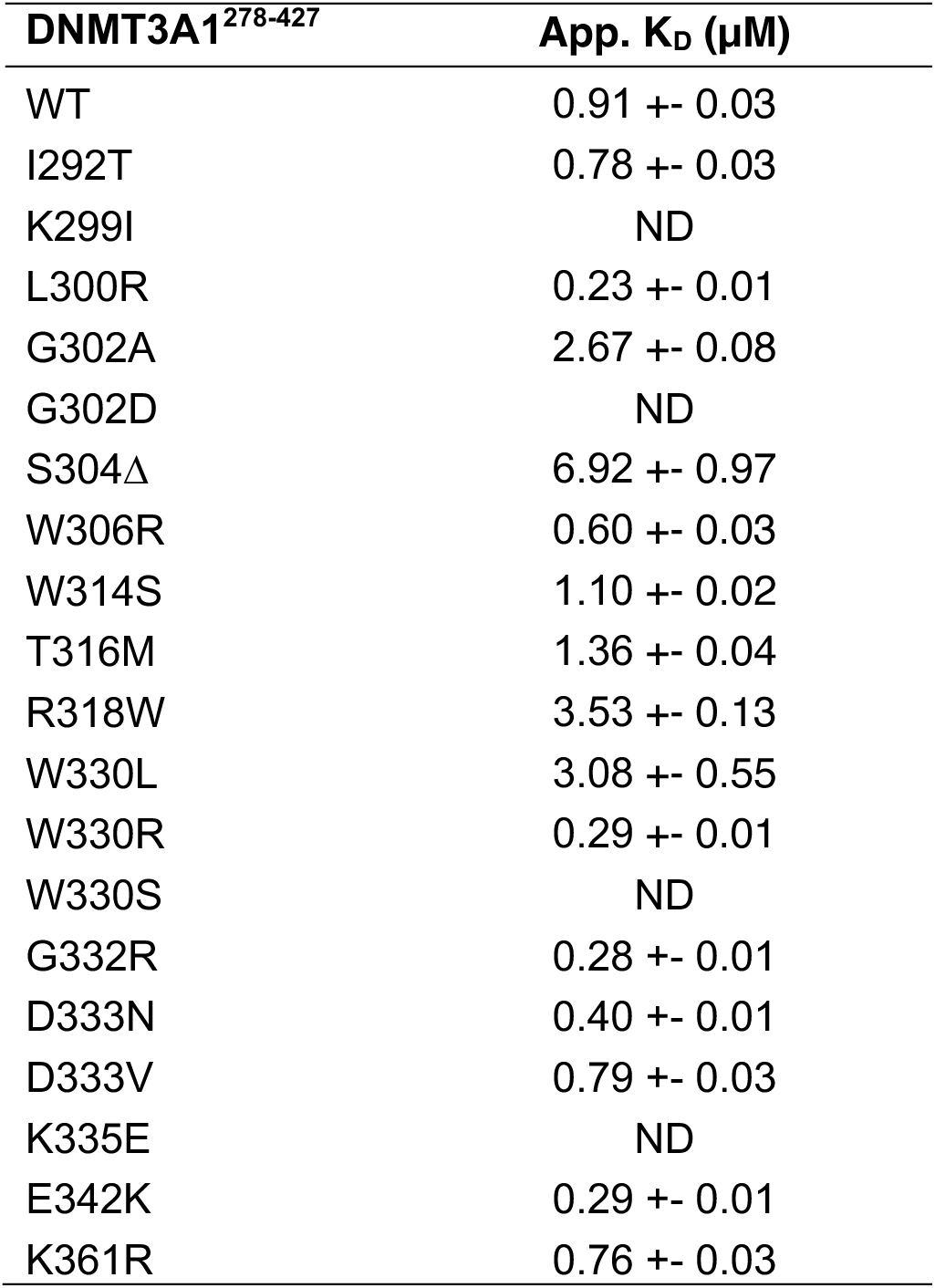
Apparent K_D_ values of DNMT3A1^278-427^ binding to 145bp DNA determined by EMSA.

DNMT3A isoforms differ in their N-terminal region and are differentially expressed during development [8, 10, 16]. DNMT3A1 contains an N-terminal region that is lacking in DNMT3A2 and contains a UDR motif that interacts with the H2AK119ub histone modification as well as with the nucleosome surface [10–13, 35]. We previously showed that a construct that contains both the PWWP domain and the DNMT3A1-specific N-terminal region (DNMT3A1^1-427^) binds better to nucleosomes than the PWWP domain alone [13]. Indeed, despite the increase of DNA binding of W330R PWWP domain we observed here (Figure 3D, Table 3), this mutation did not alter affinity for unmodified nucleosomes in the larger DNMT3A1^1-427^ fragment [13]. To test whether the additional nucleosome binding from the N-terminal region generally offsets and buffers the DNA-binding effects of PWWP mutations, we performed EMSAs with purified DNMT3A1^1-427^ WT, and HESJAS mutations S304Δ and D333N. These were incubated with unmodified and H3K36me2 modified NCPs in the presence of competitor DNA prior to separation and visualisation (Figure 4A&B; Supp. Figure 4B). Consistent with the N-terminal region making an additional, dominant contribution to nucleosome binding, DNMT3A1^1-427^ bound NCPs with an apparent 10-fold higher affinity than the isolated PWWP domain (Compare Figure 3B and Figure 4B). As found for the PWWP alone, S304Δ and D333N mutations lost preference for H3K36me2-modified nucleosomes (Figure 4B). However, neither mutation altered the affinity to unmodified NCPs, despite altered binding affinity to DNA for the PWWP domain alone (Figure 3D). This suggests that the mutant-driven DNA binding affects observed for the isolated PWWP domain are tempered in the presence of the isoform-specific N-terminal region. It therefore appears that in DNMT3A1 interactions of the N-terminal region with the nucleosome dominate and mask the charge-dependent DNA-binding effects of PWWP mutations. Because DNMT3A2 lacks this N-terminal region (Figure 1A), PWWP mutations that change DNA binding may have a greater impact on DNMT3A2 than on DNMT3A1.

**Figure 4.**
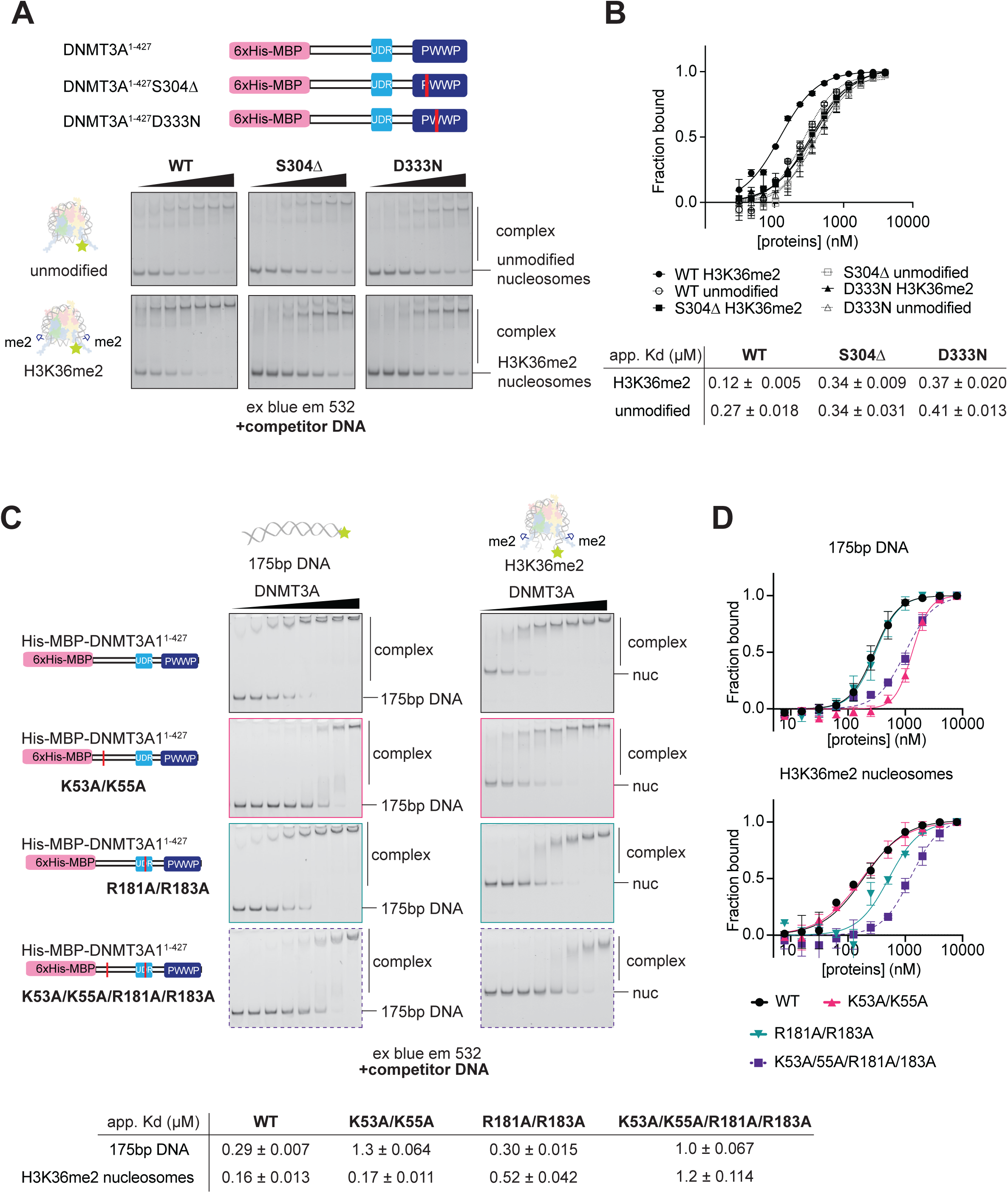
The disordered DNMT3A1 N-terminal region is involved in DNA binding and contributes to nucleosome engagement. A. EMSAs comparing the binding of DNMT3A1^1-427^ WT, S304Δ and D333N to unmodified and H3K36me2 modified nucleosomes (2.7 nM; 5′-6-FAM-labelled 145 bp Widom 601 DNA). Dilution series from, 0-4 μM performed (1.5-fold series); 156–1777 nM shown for clarity. B. Quantification of experiments A, with full concentration series from two independent experiments. C. EMSAs comparing binding of DNMT3A1^1-427^ WT, K53/55A, R181/183A or K53A/55A/R181A/183A to free 6-FAM- labelled 175bp Widom601 DNA (left, 2.3 nM) or H3K36me2 modified nucleosomes (right; 5′ 6-FAM- labelled 175bp Widom601 DNA wrapped, 2.3 nM). DNMT3A1^1-427^ WT or mutants (0-8 μM, 2x dilution series) were resolved on native PAGE gels and imaged using blue light, emission 532 nm. For clarity, 62.4-8000 nM is shown on gels. D. Quantification of C performed with full concentration series from three experiments.

### The DNMT3A1 N-terminal region contains two chromatin interacting regions

In light of the above results, we next sought to further map the interaction of the DNMT3A1 N-terminal region with nucleosomal features such as DNA and the nucleosome hostone core. The N-terminal region of DNMT3A1 contains two separable stretches of basic clusters that are intolerant to mutation (Figure 1B, light blue and black boxes). Previously, these motifs were reported to bind DNA [14, 15], with the second cluster later revised to instead contact the nucleosome surface. This is principally mediated via a negatively-charged canyon termed the acidic patch [11–13]. To test the role of the two invariant basic clusters further, we reduced the positive charge of the first region at Lys-53 and Lys-55 in DNMT3A1^1-427^ (K35A/K55A) and tested DNA binding by EMSA (Figure 4C&D; Supp. Figure 1C, Supp. Figure 4B). K35A/K55A mutation markedly reduced binding to DNA. Two neighbouring AML-associated N-terminal mutations E29A ([32, 50]) and N89S ([32]) did not appreciably affect DNA binding (Supp. Figure 4B-D), suggesting this is a specific interaction within the K53/55A motif. Despite inhibiting DNA binding, the K53/55A mutation had little effect on interaction with H3K36me2 modified nucleosomes (Figure 4C&D; Supp. Figure 1C, Supp. Figure 4B), suggesting additional N-terminal regions are the main drivers of nucleosome engagement.

Conversely, we found mutating previously characterised nucleosome contacting residues (R181A/R183A [13]) strongly reduced nucleosome binding but only marginally affected DNA binding (Figure 4C&D). Mutation of all 4 residues (K53A/K55A/R181/R183A) reduced overall DNA binding similarly to just K35A/K55A, further supporting the conclusion that the R181/183 region does not appreciably contact DNA. However, on H3K36me2-nucleosomes, the K53/55A mutation was additive to the R181A/R183A mutation reducing nucleosome binding beyond R181A/R183A alone, indicating that the two clusters concurrently contribute to nucleosome engagement. Together, these data indicate that K53/K55 contacts nucleosomal DNA, whereas R181/R183 mediates the acidic-patch interaction that is the principal driver of nucleosome affinity under these conditions. This provides the DNMT3A1 N-terminal region with multiple, partially redundant means of engaging chromatin.

### Thermal destabilisation of the isolated PWWP domain does not predict the stability of DNMT3A1 in cells

Mutations in DNMT3A give rise to a spectrum of phenotypes that correlate, in part, with their effect on protein stability. Whereas TBRS and AML mutations are frequently associated with loss of protein stability and reduced catalytic activity [26, 29, 32, 51], the gain-of-function DNA methylation of HESJAS and paraganglioma require an intact active protein [25, 27, 38]. To assess the direct effect of PWWP mutations on protein stability, we measured the thermal stability of purified DNMT3A^278-427^ and mutants by differential scanning fluorimetry (Figure 5A, Supp. Figure 5A). Mutations previously reported not to affect stability in a cellular reporter assay (I292T, G302D, W314S, T316M, E342K and K361R [32]) likewise did not alter the melting temperature (Tm) of the isolated domain. Similarly, a mutation previously shown to be destabilising in cells, G332R [32], was also destabilising in the purified domain (Figure 5A). Strikingly, L300R, but no other HESJAS mutations, markedly increased the thermal stability of the domain. As the single HESJAS patient carrying L300R exhibited the most severe phenotype [28], we hypothesised that increased stability might raise cellular protein levels and thereby amplify the gain-of-function activity.

**Figure 5.**
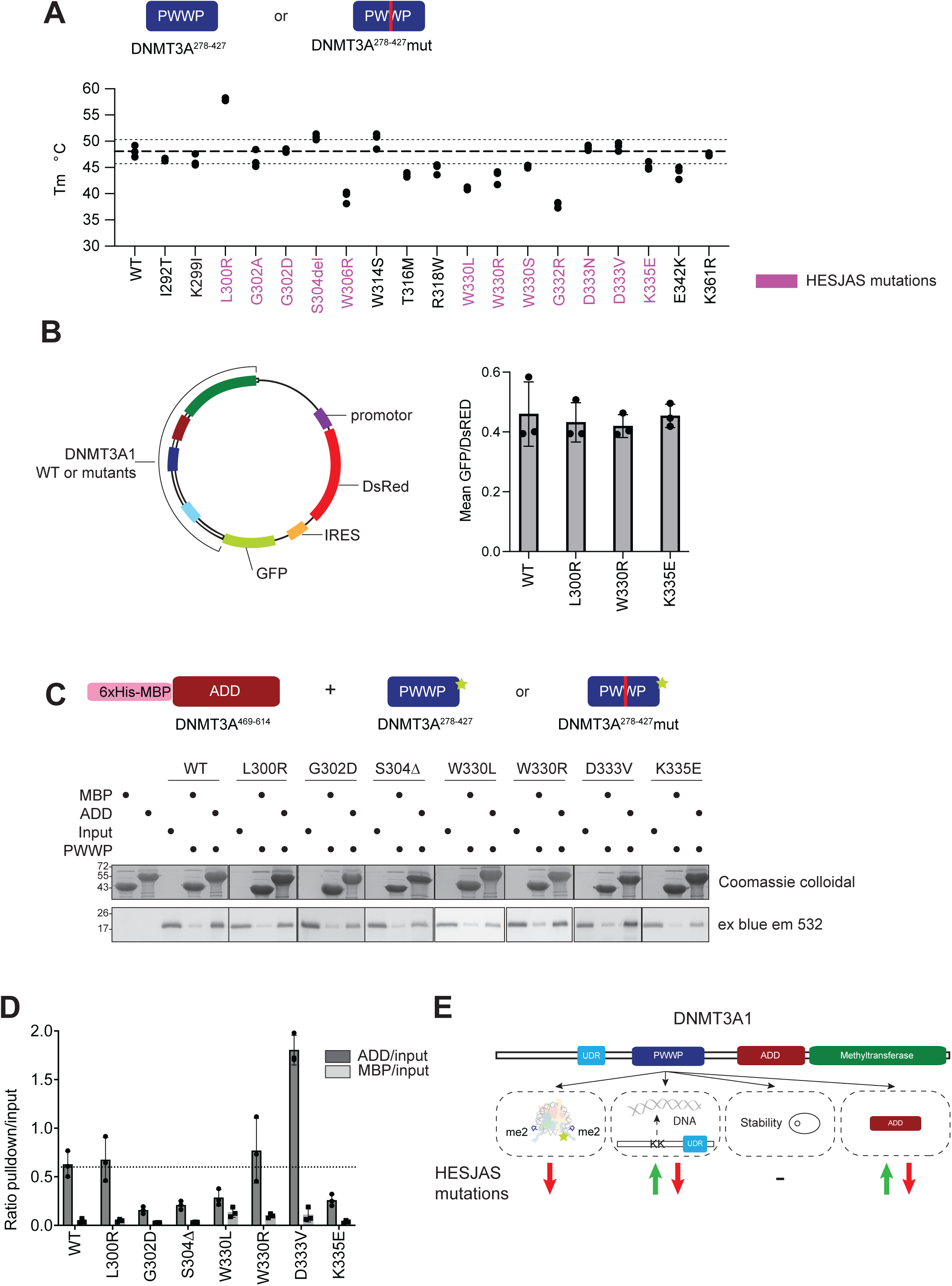
PWWP-domain mutations affect PWWP-ADD interaction, but not full-length protein stability in cells. A. Melting temperatures (Tm) of DNMT3A1^278-427^ WT and mutants (5 μM) by differential scanning fluorimetry. Tm values were determined as the peak of the first derivative of the melting curve. Three repeats. Dashed and dotted lines show the mean and 95% confidence interval for WT. HESJAS mutations shown in magenta. Full melting curves shown in Supp. Figure 5A. B. Stability of DNMT3A1 WT, L300R, W330R and K335E measured by fluorescence reporter assay. HEK293T cells were transfected with a polycistronic vector expressing DsRed and GFP-tagged DNMT3A proteins (left), reported as the mean GFP/DsRed ratio by flow cytometry. Three biological experiments performed in triplicate. Histograms showing the density distribution of GFP/dsRED ratios of one representative experiment for each mutation are shown in Supp. Figure 5B. C. Pull-down of OG488-labelled DNMT3A1^278-427^ (PWWP domain) WT and mutants by immobilised His-MBP-DNMT3A1^469-614^ (ADD domain), with His-MBP as a control. Proteins were resolved on SDS-PAGE. OG488 fluorescence (top) and colloidal Coomassie (bottom) are shown. D. Quantification of triplicate of experiments represented in C. Fluorescence of OG488- labelled DNMT3A1^278-427^ WT and mutants were quantified, expressed as the ratio of signal present in the His-MBP control lane and His-MBP-DNMT3A1^469-614^ (ADD domain) lane relative to the input. E. Schematic model summarising how PWWP-domain mutations affect H3K36me2 reading, DNA binding (charge-dependent), the PWWP-ADD interaction and protein stability, in the context of the DNMT3A1 and DNMT3A2 isoforms.

To test this, we measured the stability of full-length DNMT3A1 in cells using a bicistronic fluorescent reporter assay [32, 48], comparing WT with HESJAS mutations L300R, K335E and the previously characterised W330R [27]. K335E was selected as it does not affect stability *in vitro* and is associated with a milder phenotype [28]. DsRED and GFP-tagged DNMT3A1 WT or mutants were co-expressed in HEK293T cells, and GFP-DNMT3A1 levels were quantified by normalising intensity to individual dsRed positive cells (Figure 5B, Supp. Figure 5B). None of the mutations appreciably altered steady-state protein levels compared to the WT protein. We conclude that changes in the thermal stability of the isolated PWWP domain do not necessarily translate into altered stability of DNMT3A1 in cells. This argues that PWWP-domain mutant effects on protein stability are unlikely to be a major determinant of HESJAS severity.

### HESJAS mutations differentially affect the PWWP-ADD interaction

The catalytic activity of DNMT3A is internally regulated by other domains of the protein. The ADD domain autoinhibits the catalytic domain through a conformation that is relieved upon binding the unmodified H3 tail (H3K4me0) [3, 40, 52]. More recently, the PWWP domain has also been proposed to contribute to autoinhibition through a direct interaction with the ADD domain [40]. Consistent with this, H3K36me2 binding stimulates the activity of WT DNMT3A [13, 37, 53], which is attenuated by mutation of the PWWP that block H3K36me2 interaction [37]. Additionally, an H3K36me2-independent stimulatory effect was observed for W330R mutation on the activity of DNMT3A [40].

To test whether selected HESJAS mutations affect the PWWP-ADD interaction directly, we performed a pull-down assay using immobilised His-MBP-ADD domain (DNMT3A1^469-614^). This was incubated with WT or mutant fluorescently-labelled DNMT3A1^278-427^ (PWWP domain) and analysed by SDS-PAGE (Figure 5C, Supp. Figure 5C). Due to the low affinity of interaction [40] large amounts of both proteins were used in the assay and low levels of background binding was observed in His-MBP control. Nevertheless, a clear fluorescent-PWWP signal was retained specifically on the ADD domain pull-downs (Figure 5D). Several mutations (G302D, S304Δ, W330L and K335E) reduced the interaction with the ADD domain, but this was not a general property for all HESJAS mutations. Notably, D333V markedly increased binding, and L300R and W330R had no detectable effect. The lack of effect of W330R was unexpected, given that this mutation increases catalytic activity [40]; however, W330L did reduce binding, as previously reported for the non-disease substitution W330A [40]. This indicates that the identity of the substituting residue, and not merely loss of the wild-type residue, determines the biochemical outcome. Overall, HESJAS mutations can affect the PWWP-ADD interaction, but the effect is variable, indicating that additional factors couple this interaction to catalytic regulation.

## Discussion

In this study we used a systematic biochemical approach to dissect how mutations in the PWWP-domain associated with HESJAS, clonal haematopoiesis and paraganglioma affect the multiple functions of DNMT3A (Figure 5E).

There is a clear correlation between HESJAS and loss of H3K36me2 binding: we observed that every HESJAS mutation in the purified PWWP domain abolished binding to H3K36me2 modified nucleosomes, whereas clonal haematopoiesis and paraganglioma mutations, with one exception, did not. The exception, K299I, also ablated H3K36me2 binding and produces a HESJAS-like chromatin gain-of-function DNA methylation phenotype [25]. As a somatic mutation, its lack of developmental phenotype is expected. Consistent with a shared mechanism, HESJAS patients have been reported to develop paragangliomas [27, 28], suggesting that loss of H3K36me2 binding and associated gain-of-function may drive paraganglioma with or without HESJAS. In contrast, the same-residue K299N mutation causes AML and is predicted to impair DNA binding and catalytic activity [32, 37]. The germline paraganglioma mutation R318W had a minimal effect within the error of the assay on H3K36me2 binding but reduced DNA binding [25]. Thus, while HESJAS is tightly linked to germline mutations that abolish H3K36me2 recognition, DNMT3A-linked paraganglioma appears to arise through several distinct mechanisms. Further investigations are required to clarify whether this involves H3K36me2 and/or DNA binding in concert with other mutations and functional alterations.

The PWWP domain of DNMT3A binds DNA [7, 39], analogous to the PWWP domains of LEDGF [45, 46] and DNMT3B [42, 48]. We show that disease mutations cluster on the predicted interface between the PWWP domain and the nucleosome (Figure 1C) and can alter the interaction with DNA largely through changes in charge (Figure 3D). Indeed, W330R is predicted to be internally buried in the methyl-lysine binding aromatic cage but is still sufficiently solvent exposed to affect DNA interaction potential. Because W330R and E342K increase both DNA binding and catalytic activity independently of H3K36me2 [37], PWWP-domain DNA binding may be mechanistically linked to catalytic output, but may also be connected to a relief in autoinhibitory activity. Importantly, however, inclusion of the disordered N-terminal region of DNMT3A1 overcame the effects of mutations on nucleosome binding *in vitro*, most likely through its additional contacts with the nucleosome surface and DNA. This region is critical for chromatin engagement through residues R181/183, which interact with the nucleosome acidic patch [10, 13] and contains a DNA-binding motif at K53/K55 (this study, [14, 15]). Here, we show that the nucleosome surface interaction is essential for affinity, but the DNA interaction of K53/55 is additive to this interaction on nucleosomes. Despite the overall disorder of the N-terminal region, AlphaMissense predicts the K53/55 motif to be pathogenicity-sensitive (Figure 1B), suggesting this additional DNA interaction could be physiologically relevant. As DNMT3A2 lacks the N-terminal region (Figure 1A), PWWP mutations that alter DNA binding may have a more pronounced effect on DNMT3A2 than DNMT3A1. Such mutations might therefore make isoform-specific perturbations during the developmental window where DNMT3A2 predominates, leading to specific developmental disorder phenotypes [8, 10, 16].

The PWWP domain has been proposed to contribute to autoinhibition of the catalytic domain via the ADD domain [40], as also suggested for the paralogue DNMT3B [41, 42]. The non-disease substitution W330A disrupts the PWWP-ADD(-catalytic) domain interaction [41]. In addition, HESJAS mutations W330R and D333N stimulate DNMT3A activity on DNA in the absence of H3K36me2 [40], suggesting that disruption of the PWWP-ADD interaction could promote aberrant hypermethylation seen in HESJAS. We were therefore surprised that different HESJAS mutations had divergent effects on the PWWP-ADD interaction (Figure 5D), most notably the lack of altered binding in pull-down assay of W330R. These differences may reflect the different protein constructs and methods used here and elsewhere [40]. A mutation-driven effect may also be offset by intrinsic flexibility of the PWWP and ADD domains [54]: the ADD domain adopts different conformations in crystal and cryo-EM structures [3, 40], and the PWWP domain is poorly resolved or absent in cryo-EM structures of DNMT3A [13, 55–58], indicating considerable conformational heterogeneity. In addition, modulation of the PWWP-ADD interaction may not be unique to HESJAS: the clonal haematopoiesis mutation E342K also stimulates activity [37]. All mutations that increased catalytic activity also increased DNA binding (W330R, D333N and E342K in this study and [37]; W330A and the ADD mutation R544A in [40]). Conversely, all mutations that reduced the PWWP-ADD interaction in our assay (G302D, S304Δ, W330L and K335E; Figure 5D) also reduced DNA binding (Figure 3D). This raises the possibility that PWWP-domain DNA binding and the PWWP-ADD interaction are mechanistically coupled, although further work will be needed to define how the ADD and PWWP domains jointly regulate the catalytic domain on chromatin.

Our data reinforce the emerging view that PWWP-domain mutations are functionally pleiotropic and that even different substitutions at a single residue can have distinct consequences. This is exemplified by substitutions of tryptophan at position 330: W330R, W330L and W330A each produce different effects on H3K36me2 binding, DNA binding and the PWWP-ADD interaction (this study; [37, 40]). Such residue- and substitution-specific behaviour underscores the value of characterising individual clinical variants rather than inferring function from the wild-type residue alone, and helps explain the range of phenotypes and clinical severities observed across PWWP-domain mutations. Even subtle changes such as G302A were sufficient to both reduce DNA and H3K36me2-nucleosome binding, without globally affecting PWWP stability. Alanine is less flexible than glycine and G302 is highly kinked in a loop adjacent to the H3K36me2-binding pocket, suggesting that the PWWP binding interfaces are disrupted even with this minor alteration.

Our results carry two broader caveats necessitated by the reductionist approach. There is a large conceptual difference between *in vitro* biochemistry and the higher complexity inside of cells and further still to whole organismal phenotypes. Firstly, effects observed with isolated domains may not reflect the behaviour of the full-length protein. For example, the DNA-binding effects of PWWP mutations were largely masked in the DNMT3A1^1-427^ construct, and changes in the thermal stability of the isolated PWWP domain did not translate into altered stability of full-length DNMT3A1 in cells. Interpretations based on single-domain constructs should therefore be made with care. Indeed, even the DNMT3A1^1-427^ construct is missing other chromatin and DNA binding capabilities compared to the full-length protein. Second, DNMT3A has multiple reported binding partners [24, 58–61]. Indeed, DNMT3A functions as a hetero-oligomer and disease mutations are frequently heterozygous. As a result, the interplay between wild-type and mutant protein within a complex and the potential for dominant-negative or dominant-gain effects, as described for R882H [31, 62], remains an important open question that our biochemistry does not address.

The PWWP domain of DNMT3A has been studied chiefly for its role in reading H3K36me2, which is essential for correct genomic targeting and embryonic development. Our results add to a growing appreciation that the domain is multifunctional: contributing to DNA binding and to autoinhibition of catalytic activity. We show that disease mutations affect these functions in distinct, sometimes amino acid-specific ways, but a unifying observation is that all tested HESJAS mutations ablate H3K36me2 interaction. This implies that loss of this interaction is the key molecular defect for pathogenesis of HESJAS. This work builds additional biochemical insight into understanding how PWWP-domain mutations give rise to the divergent phenotypes of HESJAS, paraganglioma and clonal haematopoiesis.

## Methods

### Predicted pathogenicity of DNMT3A1 amino acid mutations

Per-residue missense pathogenicity scores for DNMT3A1 were obtained from the AlphaMissense database [43]. For each residue, scores across all 19 possible amino-acid substitutions were summed to give a cumulative pathogenicity score, which was plotted against residue number (Figure 1B)

### AlphaFold prediction

A prediction of the structure of DNMT3A1^278-427^ bound to an H3K36me2 modified nucleosome wrapped with 175 bp Widom 601 DNA was made using AlphaFold 3 [44]. For clarity, the histone tails were truncated in Figure 1C, shown are H2A 10-123, H4 13-103, H2B 27-126 and H3 26-136. Full histone tails are shown in Supp. Figure 1A.

### Generation of plasmids

A list of constructs can be found in Supp. Table 2. Bacterial expression constructs for DNMT3A were generated from cDNA (Addgene #35521, #35523) by ligation independent cloning (LIC) into a pLIC-His6-MBP-TEV vector. DNMT3A1^278-427^ and DNMT1^1-427^ point mutants were generated by site-directed mutagenesis of the corresponding wild-type constructs [13]. DNMT3A1 constructs for fluorescent protein stability assays were created using Gibson assembly [63]. Histone constructs were described previously [64–66]. Polycistronic vector constructs expressing DsRed and GFP-tagged DNMT3A1 proteins (WT or mutant) were generated by cloning DNMT3A variants into the pLenti-DsRed-IRES-EGFP vector (Addgene plasmid #92194; gift from Huda Zoghbi).

### Protein purification

#### Expression and purification of wild type and mutant DNMT3A1^278-427^

His6-MBP-tagged DNMT3A1^278-427^ constructs were expressed and purified as described before [13] with minor adjustments. In short, His-MBP tagged constructs were expressed in BL21 (DE3 RIL) E. coli cells with 400 μM IPTG at 18 °C overnight in 2xTY medium (16 g/l tryptone, 10 g/l yeast extract, 5 g/l NaCl, pH 7.0). Cell pellets were resuspended in in lysis buffer (25 mM sodium phosphate pH 7.5, 400 mM NaCl, 0.1% (v/v) Triton, 10% (v/v) glycerol, 1 mM TCEP, 1 mM AEBSF, 1X protease inhibitor cocktail (2.2 mM PMSF, 2 mM benzamidine HCl, 2 μM leupeptin, 1 μg/ml pepstatin A), 4 mM MgCl2, 5 μg.mL−1 DNAse and 500 μg/ml lysozyme) and stirred at 4 °C before additional lysis using a sonicator (Branson Digital Sonifier®, 1/8th inch microtip, 2 sec on, 2 sec off for total 20 s per 3 L of the original culture at 50% amplitude). Lysate was cleared by centrifugation at 39,000 × g for 25 min and the supernatant was filtered through a 0.45 μm filter. Lysate was then loaded on Ni-NTA beads (Qiagen, 1 ml per litre culture), washed with 25 column volumes (CV) 15 mM sodium phosphate, 500 mM NaCl, 10% glycerol, 15 mM imidazole, 1mM TCEP and eluted with 5CV 20 mM Tris pH 7.5, 400 mM NaCl, 300 mM imidazole, 10% glycerol, 1mM TCEP. The His-MBP tag was then cleaved using TEV protease (1:30 TEV:protein w/w) while dialysing to 15 mM HEPES pH7.5, 150 mM NaCl, 5% glycerol, 1 mM DTT, 4 mM sodium citrate overnight at 4 °C and run over a Ni-NTA column (chelating HP conjugated with NiSO4, Cytiva) to remove the tag. The cleaved protein was further purified using size-exclusion chromatography (HiLoad 16/600 Superdex 75 pg, Cytiva, equilibrated in 15 mM HEPES pH7.5, 150 mM NaCl, 5% glycerol, 1 mM DTT), frozen in liquid nitrogen and stored at -80 °C. Concentration was determined by A₂₈₀ (NanoDrop); purity was assessed by SDS-PAGE (0.5 and 1 μg per lane; Figure 1D).

Purified DNMT3A1^278-427^ WT and mutants were analysed by intact-protein LC-MS to confirm identity. Measured masses agreed with calculated values for all constructs (Supp. Table 1).

#### Expression and purification of wild type and mutant DNMT3A1^1-427^

DNMT3A1^1-427^ wild type and mutants were purified as previously described previously [13], in short: His-MBP tagged constructs were expressed in BL21 (DE3 RIL) E. coli cells with 400 μM IPTG at 18 °C overnight in 2xTY medium (16 g/l tryptone, 10 g/l yeast extract, 5 g/l NaCl, pH 7.0). Cells were harvested at 4000 x g for 10 min and pellets were resuspended in in lysis buffer (25 mM sodium phosphate pH 7.5, 400 mM NaCl, 0.1% (v/v) Triton, 10% (v/v) glycerol, 2 mM β-mercaptoethanol, 1 mM AEBSF, 1X protease inhibitor cocktail (2.2 mM PMSF, 2 mM benzamidine HCl, 2 μM leupeptin, 1 μg/ml pepstatin A), 4 mM MgCl2, 5 μg.mL−1 DNAse and 500 μg/ml lysozyme) and stirred at 4 °C for an hour before additional lysis using a sonicator (Branson Digital Sonifier®, 1/8th inch microtip, 2 sec on, 2 sec off for a total 20 s per 3 L of the original culture at 50% amplitude). Lysate was cleared by centrifugation at 39,000 × g for 25 min and the supernatant was filtered through a 0.45 μm filter. Lysate was then loaded on Ni-NTA beads (Qiagen, 1 ml per litre culture), washed with 25 column volumes (CV) 15 mM sodium phosphate, 500 mM NaCl, 10% glycerol, 15 mM imidazole, 2mM β-mercaptoethanol and eluted with 5CV 20 mM Tris pH 7.5, 400 mM NaCl, 300 mM imidazole, 10% glycerol, 2mM β-mercaptoethanol. The eluted protein was concentrate and diluted to 100 mM NaCl using 20 mM HEPES pH 7.5, 10% glycerol, 1 mM DTT and loaded on 5 mL HiTrap Q HP cation exchange chromatography column (Cytiva), washed with 15 CV 20 mM HEPES pH 7.5, 100 mM NaCl,10% glycerol, 1 mM DTT and eluted with a salt gradient (0–50% 20 mM HEPES pH 7.5, 1 M NaCl, 10% glycerol 1 mM DTT) and further purified on a HiLoad Superdex 200 16/600 (Cytiva) equilibrated with 15 mM HEPES pH 7.5, 150 mM NaCl, 1 mM DTT, 5% (v/v) glycerol. Fractions containing pure protein were pooled and concentrated in a 30 kDa MWCO centrifugal filter. Proteins were flash frozen in liquid nitrogen and stored at −80 °C.

### Formation of 145 bp and 175bp nucleosomes

Unmodified and H3K36me2 modified nucleosomes were prepared as described [13]. In short:

#### Histone purification

Histones were expressed in E. coli (BL21 (DE3 RIL)), resolubilised from inclusion bodies, purified by cation exchange and dialysed into 1mM acetic acid before lyophilisation and stored at -20 °C.

#### Native chemical ligation

Native chemical ligation of H3K36me2 histones was done as described in [13], based on [67, 68]. H3 Δ1-44 T45C C110A histone was reacted with H3 1-43 K36me 2 S-Bnzl peptide (Peptide Synthetics), dialysed into 7 M Urea, 25 mM Tris pH 7.5, 20 mM NaCl, 1 mM EDTA, 2 mM β-mercaptoethanol and purified using cation exchange chromatography.

#### Octamer refolding

Lyophilised histones were resuspended in 20 mM Tris pH 7.5, 6 M guanidine, 10 mM DTT and combined in a ratio of 1:1:1.5:1.5 H3, H4, H2A, H2B, dialysed to 15 mM Tris pH 7.5, 2 M NaCl, 5 mM β-mercaptoethanol, 1 mM EDTA and purified using a HiLoad Superdex 200 16/600 (> 10 mg total protein, Cytiva) or Superdex 200 Increase 10/300 GL (<10 mg total protein, Cytiva) in 15 mM Tris pH 7.5, 2 M NaCl, 1 mM EDTA, 5 mM β-mercaptoethanol. Octamer fractions were pooled, concentrated and stored in 50% (v/v) glycerol at −20 °C.

#### PCR amplification of 145 and 175 bp Nucleosome DNA

DNA fragments (Widom 601 145bp and 175 bp DNA) were generated by PCR amplification followed by ion exchange chromatography (ResourceQ column, Cytiva). Pure fractions were concentrated by ethanol precipitation and resuspended in 10 mM Tris pH8.

#### Nucleosome wrapping

Octamers were combined with DNA in a 1:1.2 ratio and dialysed in a salt reduction gradient (2 M KCl to 0.2 M KCl in 15 mM HEPES pH 7.5, 1 mM DTT, 1 mM EDTA). Nucleosomes were dialysed to 15 mM HEPES pH 7.5, 100 mM NaCl, 1 mM DTT, 1 mM AEBSF, purified by PEG precipitation and resuspended in 15 mM HEPES pH 7.5, 100 mM NaCl, 1 mM DTT, 1 mM AEBSF.

### Fluorescence polarisation assays

Nucleosomes wrapped with 6-carboxyfluorescein (5′ 6-FAM) labelled 145bp Widom601 DNA (6.7 nM) were mixed with DNMT3A1^278-427^ WT and mutants (50-0 µM 2 x dilution series, except W330S, G332R and K299I 0-25 µM due to limited concentration) in FP buffer (15 mM HEPES pH7.5, 150 mM NaCl, 0.005% (v/v) NP-40, 0.05 mg/mL BSA, 5% (v/v) glycerol, 1 mM DTT) in a final volume of 25 µL. Samples were incubated for 15 min at room temperature and fluorescence polarisation measured with a SpectraMax iD5 (Molecular Devices) plate reader with 480 nm excitation and 540 nm emission polarised filters (cut-off at 530 nm). Fluorescence polarisation was calculated using

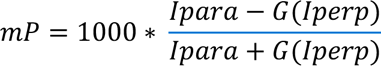

Where mP = fluorescence polarisation, Ipara = parallel fluorescence intensity values, Iperp = perpendicular fluorescence intensity values, G = G factor.

Results were baseline subtracted (no protein) and plotted against the concentration of protein. Binding curves were analysed in GraphPad Prism 11 using non-linear regression – one site -- specific binding. Independent experiments were performed 3 times in technical duplicates on a plate, except W330S which was performed twice in duplicate due to restricted amount of protein.

### Electrophoretic mobility shift assays

#### DNMT3A1^278-427^ and DNMT3A1^1-427^ with nucleosomes

Unmodified or H3K36me2 modified nucleosomes wrapped with 6-carboxyfluorescein (5′ 6-FAM) labelled 145bp Widom601 DNA (2.7 nM) were incubated with a 1.5 x dilution series of DNMT3A1^278-427^ WT and mutants (0-25 μM) or DNMT3A1^1-427^ WT and mutants (0-4 μM) in EMSA buffer 1 (15 mM HEPES pH7.5, 75 mM NaCl, 0.05% (v/v) Triton X-100, 0.05 mg/mL BSA, 10% (v/v) glycerol, 1 mM DTT, 0.5 mg/mL salmon sperm DNA (low molecular weight 31149-10g-f, Sigma-Aldrich), 8% (w/v) sucrose, 0.01% (w/v) bromophenol blue) in 12 μl. Samples were incubated on ice for 1 hour and products were separated on 5% 19:1 acrylamide native PAGE gels using 1xTris Glycine as running buffer at 100V for 90 min at 4°C. Gels were imaged for FAM signal (Excitation Blue light, Emission 532nm) using Bio-Rad ChemiDoc MP. Quantified using Image Lab (Bio-Rad) and binding curves were analysed in GraphPad Prism 10 using non-linear regression (specific binding with hill slope).

#### DNMT3A1^278-427^ with 145 bp DNA

6-carboxyfluorescein (5′ 6-FAM) labelled 145 bp Widom601 DNA (2.7 nM) was mixed with 1.5 x dilution series with DNMT3A1^278-427^ WT or mutants (0-12.5 μM) in EMSA buffer 2 (15 mM HEPES pH 7.5, 75 mM NaCl, 0.05% (v/v) TritonX100, 0.05 mg/mL BSA, 10% (v/v) glycerol, 1 mM DTT, 8% (w/v) sucrose, 0.01% (w/v) bromophenol blue) in 12 μl. Samples were incubated on ice for 1 h and separated, imaged and quantified as above.

#### DNMT3A1^1-427^ K53/55A, R181/183A and K53/55A R181/183A

Nucleosomes wrapped with 5′-6-FAM-labelled 175 bp Widom 601 DNA, or free 5′-6-FAM-labelled 175 bp DNA (2.3 nM), were incubated with a two-fold dilution series of protein (0-8 μM) in EMSA buffer 1 (with salmon-sperm DNA) and processed as above.

#### DNMT3A1^1-427^ E29A and N89S

5′-6-FAM-labelled 40 bp DNA (10 nM) was incubated with a 1.5-fold dilution series of DNMT3A1₁₋₄₂₇ WT, E29A or N89S (0-2.5 μM) in EMSA buffer 2 (15 μL final volume). After 30 min at room temperature, complexes were resolved (100 V, 45 min) and processed as above.

### Thermal stability assay (Differential Scanning Fluorimetry)

DNMT3A1^278-427^ WT and mutants (5 µM) were mixed with 5x SYPRO orange (Life Technologies) in 15 mM HEPEs pH 7.5, 150 mM NaCl, 1mM DTT, 0.25 % glycerol in a total volume of 50 µL. The proteins were heated from 25 °C to 75 °C degrees with 0.5 degree 0.5 degree intervals on a RT-PCR device (Biometra TOptical, excitation/emission λ = 490/580 nm) and the fluorescence intensity was measured. The fluorescence was normalized and plotted against the temperature. The inflection point of the curve Tm was determined by taking the maximum of the first derivative of the curves. Three independent replicates were performed.

### Cellular fluorescent protein stability assay

HEK293T cells were transfected with a polycistronic vector expressing DsRed and GFP-tagged DNMT3A1 proteins (WT or mutant). Cells transfected with plasmids expressing GFP alone, DsRed alone, or no plasmid were used as controls. 48 hours after transfection, cells were analysed by flow cytometry on BD FACSAria II. Matrix compensation was set-up using negative (no plasmid), DsRed only and GFP only cells. Each DNMT3A mutant was analysed in technical triplicate within each experiment, and three independent transfection experiments were performed.

Flow cytometry data were analysed using FCS Express (De Novo Software). GFP and DsRed fluorescence intensities were exported for individual DsRed-positive cells, and the GFP/DsRed fluorescence ratio was calculated for each cell. The mean fluorescence ratio from the technical replicates was calculated for each experiment, and the mean value across the three independent transfection experiments was subsequently determined.

### Pull-down assays

DNMT3A1^278-427^ WT and mutants were labelled with Oregon green 488 succinimidyl ester (AAT Bioquest, Inc) at a 2:1 dye:protein molar ratio for 3 hours in 20 mM HEPES pH 8.5, 150 mM NaCl, 5% glycerol, 1 mM DTT. Free dye was removed by dialysis in 10 kD MWCO Slide-A-Lyser MINI Dialysis Units (Thermo Scientific) into pull-down buffer (20 mM Tris pH7.5, 120 mM KCl, 0.01%NP40, 15 mM imidazole, 10% glycerol, 100 μg/mL BSA, 1 mM AEBSF, 1 mM beta-mercaptoethanol). Proteins were flash frozen in liquid nitrogen and stored at -80 degrees.

His-MBP and His-MBP-DNMT3A^469-614^ (ADD domain) were expressed in *E. coli* BL21 (DE3 RIL) cells with 400 μM IPTG at 18 °C overnight in 200 mL 2xTY medium. Cell pellets were resuspended in in lysis buffer (5 mL lysis buffer per 1 gram of pellet, 25 mM sodium phosphate pH 7.5, 400 mM NaCl, 0.1% (v/v) Triton, 10% (v/v) glycerol, 2 mM beta-mercaptoethanol, 1 mM AEBSF, 1X protease inhibitor cocktail (2.2 mM PMSF, 2 mM benzamidine HCl, 2 μM leupeptin, 1 μg/ml pepstatin A), 4 mM MgCl2, 5 μg.mL−1 DNAse and 500 μg/ml lysozyme) and stirred at 4 °C for one hour before additional lysis using a sonicator (2 s on, 2 s off for total of 8 seconds per 6 mL lysate at 50% amplitude). Lysate was cleared by centrifugation at 17,000 × g for 10 min, the volume was equalised between conditions and 15 mM imidazole was added. 100 μL cOmplete™ His-Tag Purification Resin (Roche, 5893682001) per 100 mL expressed culture was added and incubated for 2 hours rotating at 4 °C. Beads were washed with 4x 1mL wash buffer (15 mM Tris pH7.5, 400 mM NaCl, 10% glycerol, 2 mM beta-mercaptoethanol, 15 mM imidazole) and another 3x 1 mL pull-down buffer (20 mM Tris pH7.5, 120 mM KCl, 0.01%NP40, 15 mM imidazole, 10% glycerol, 100 µg/mL BSA, 1 mM AEBSF, 1 mM beta-mercaptoethanol). Beads were used directly or 100 µL of pull-down buffer was added and stored overnight at 4 °C.

20 µg of Oregon Green 488 labelled DNMT3A1^278-427^ WT or mutants was incubated with 20 µL beads containing His-MBP or His-MBP-ADD in a total volume of 100 µL for 2 hours at 4 °C. Beads were washed three times, and bound protein was eluted in 2× SDS loading buffer, resolved on 17% SDS-PAGE and imaged using excitation blue and emission 532nm followed by staining with colloidal Coomassie stain.

## Data Availability

All raw and processed data is available to for sharing via the corresponding author.

## Declaration of generative AI use

Claude Opus 4.8 a large language model available via the University of Edinburgh ELM GPT architecture was used to identify typographical errors and reducing the word count of the manuscript. The authors remain fully responsible for the content and conclusions of the manuscript.

## Acknowledgements

MDW’s work is supported by a Sir Henry Dale Fellowship from the Wellcome Trust (210493/Z/18/Z) and Medical Research Council (T029471/1). WR is funded by the Wellcome trust integrative cellular mechanisms PhD program (218470). DS and FT’s work is supported by an MRC grant (UKRI576). This work was supported by the Edinburgh Protein Production Facility (EPPF). FACS data were generated by Michael Rennie in the Institute of Genetics and Cancer FACS facility. We thank Van Kelly in SIRCAMS School of Chemistry, University of Edinburgh for intact mass spec analysis. We thank Andrew Jackson for discussions and sharing information on HESJAS mutations. For the purpose of open access, the author has applied a Creative Commons Attribution (CC BY) licence to any Author Accepted Manuscript version arising from this submission.

## Author contributions

MDW conceived the study and supervised the project. HW and MDW designed the experiments (unless otherwise stated), analysed the data, and wrote the manuscript, with input from the other authors. GC, YZ, WR, FM and HW purified protein and DNA components. Biochemical assays were performed by GC, FM and HW. FT and DS performed mammalian cell reporter assays.

## Supplementary Figure legends

**Supplementary Figure 1. Predicted model validation, purified proteins and nucleosome reagents.**

A. All 5 model predictions generated by AlphaFold 3 of DNMT3A1^278-427^ with an H3K36me2 modified nucleosome (175 bp Widom 601 DNA), coloured by predicted local distance difference test (pLDDT) confidence score.

B. Predicted aligned error (PAE) plots for the 5 models shown in A

C. Native and SDS PAGE gel analysis of DNA and nucleosomes reagents used in this study. Asterisk indicates ripped gel for DNA load

**Supplementary Figure 2. Fluorescent polarisation assays of DNMT3A1^278-427^ mutants with H3K36me2 modified nucleosomes.**

Fluorescent polarisation assays of DNMT3A1^278-427^ mutants binding to H3K36me2 modified nucleosomes. DNMT3A1^278-427^ WT or mutants (0-50 μM, 2x dilution series) were incubated with H3K36me2 modified nucleosomes wrapped with 5′ 6-FAM-labelled 145bp Widom601 DNA (6.7 nM) for 15 min at room temperature. Fluorescence polarisation was measured at 480 nm excitation and 540 nm emission. 3 independent experiments with technical duplicate per assay, except K299I and W330S (N = 2). K_D_ values are shown in Table 3. E342K showed a similar affinity to WT, but overall a higher ceiling for fluorescence anisotropy. HESJAS mutations marked with magenta box.

**Supplementary Figure 3. Representative EMSA gels for all DNMT3A1^278-427^ mutants binding to DNA.**

Representative EMSA gels analysing binding of DNMT3A1^278-427^ WT and mutants to 5′ 6- FAM- labelled 145bp Widom601 DNA. Proteins were incubated with DNA at a concentration of 0-12.5 μM, 1.5 x dilution series with the exception of G302D and K335E done with 2 x dilution series. One repeat of L300R was done at 0-750 nM, 1.5x dilution series to cover lower concentrations. HESJAS mutations marked with magenta box.

**Supplementary Figure 4. DNA binding of DNMT3A1^278-427^ and DNMT3A1^1-427^ constructs**

A. Quantification of EMSAs (Supp. Figure 3) of DNMT3A1^278-427^ WT or mutations to 5′ 6- FAM- labelled 145bp Widom601 DNA. Experiments repeated 2 or 3 times, except W330S, which was performed once due to limited amount of protein. HESJAS mutations marked with magenta box.
B. SDS PAGE analysis of purified 6xHis-MBP-TEV-DNMT3A1^1-427^ WT and mutants used in this study
C. EMSAs comparing binding of DNMT3A1^1-427^ WT, E29A or N89S to 5′ 6-FAM- labelled 40 bp DNA. DNMT3A1^1-427^ WT, E29A or N89S (0-1500 nM, 1.5x dilution series) were incubated with 6-5’FAM labelled 40bp DNA (10 nM).
D. Quantifications of experiments done in B. with full concentration series from three experiments.

**Supplementary Figure 5. The effect of disease mutations on the stability of DNMT3A1**

A. Differential scanning fluorimetry melting curves for DNMT3A1^278-427^ WT and mutants. Data was normalised to baseline and 3 mean of three replicates plotted. HESJAS mutations marked with magenta box.
B. Representative single-cell GFP/DsRed ratio intensity distributions for full-length DNMT3A1 WT, L300R, W330R and K335E measured by fluorescence reporter assay. GFP-tagged DNMT3A1 WT or mutants were expressed polycistronically with dsRED and analysed by flow cytometry.
C. SDS PAGE analysis of DNMT3A1^278-427^ WT and mutants after labelling with OG-488. Fluorescence was measured using excitation blue light emission 532 nm followed by staining with colloidal Coomassie stain.

## Supplementary Tables

**Supplementary Table 1.** Intact Mass spectrometry of purified DNMT3A1^278-427^ WT and mutants.

| **DNMT3A1<sup>278-427</sup>** | **calculated MW (Da)** | **measured mass (m/z)** |
| --- | --- | --- |
| WT | 17098.33 | 17098.35 |
| I292T | 17086.27 | 17085.85 |
| K299I | 17083.31 | 17083.35 |
| L300R | 17141.35 | 17141.36 |
| G302A | 17112.35 | 17112.41 |
| G302D | 17156.36 | 17156.38 |
| S304Δ | 17011.25 | 17011.2 |
| W306R | 17068.3 | 17068.28 |
| W314S | 16999.19 | 16999.16 |
| T316M | 17128.41 | 17128.44 |
| R318W | 17128.35 | 17128.35 |
| W330L | 17025.27 | 17025.29 |
| W330R | 17068.30 | 17068.31 |
| W330S | 16999.19 | 16999.17 |
| G332R | 17197.46 | 17197.45 |
| D333N | 17097.34 | 17097.33 |
| D333V | 17082.37 | 17082.34 |
| K335E | 17099.27 | 17099.28 |
| E342K | 17097.38 | 17097.38 |
| K361R | 17126.34 | 17126.42 |

**Supplementary Table 2:**
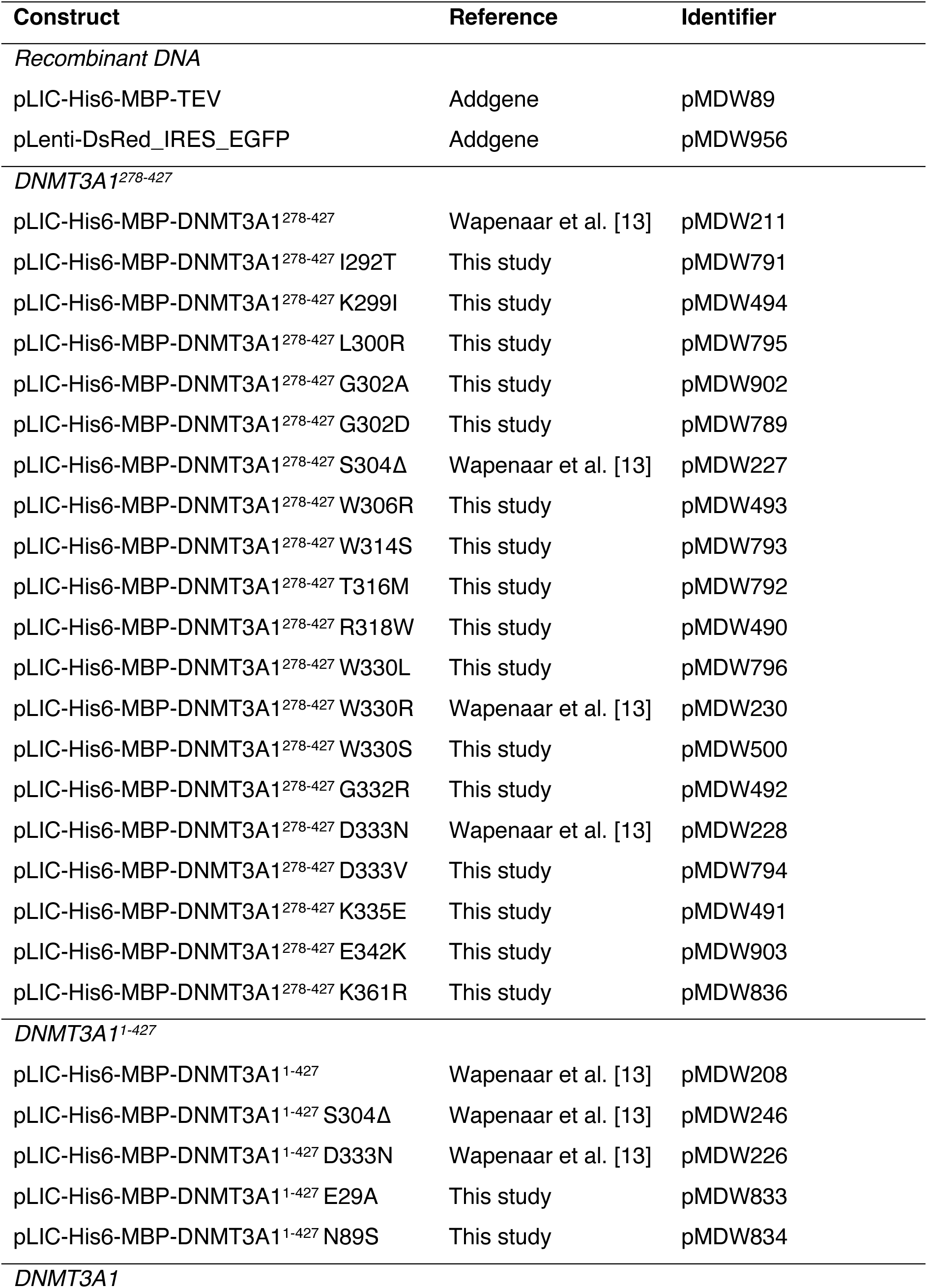
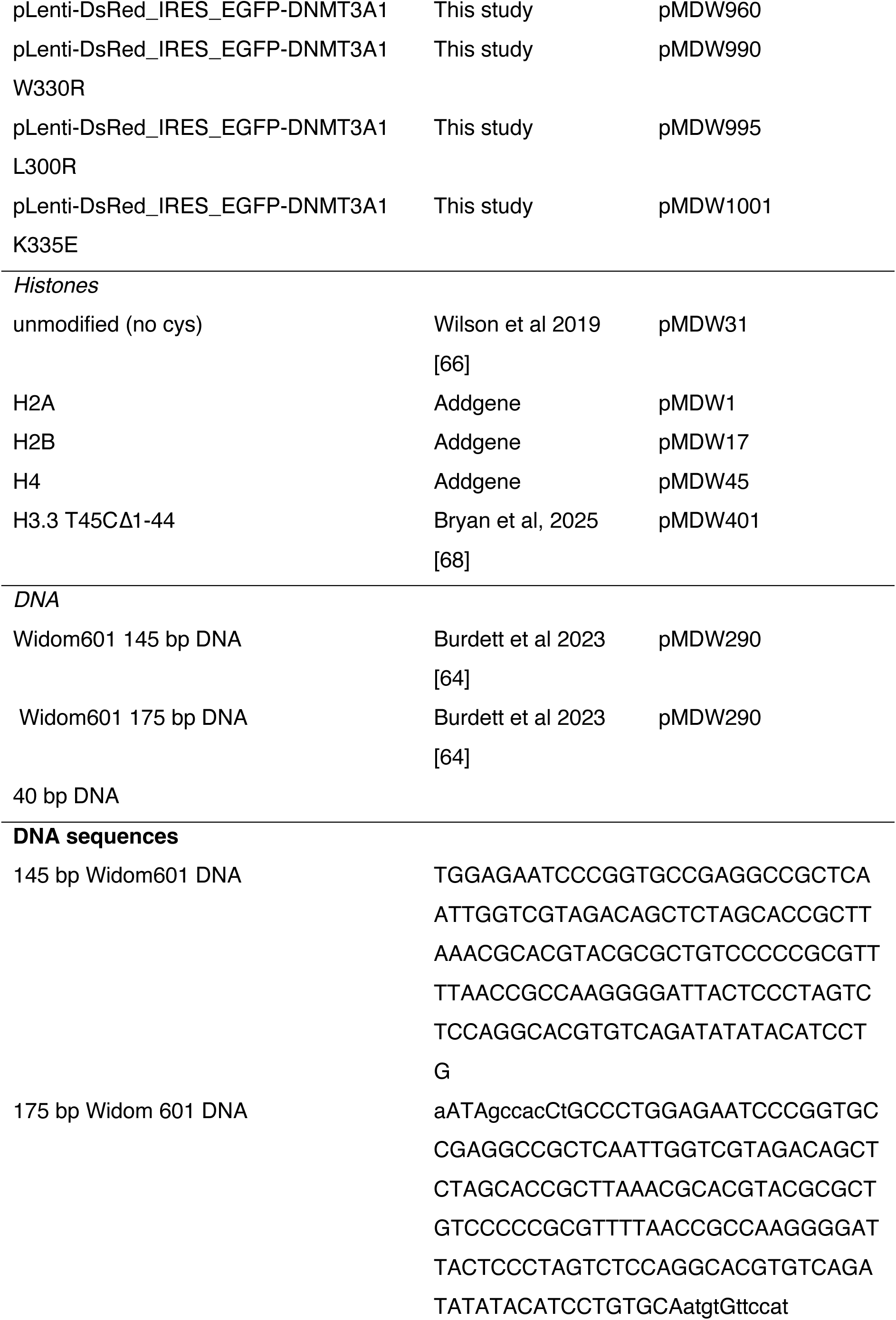
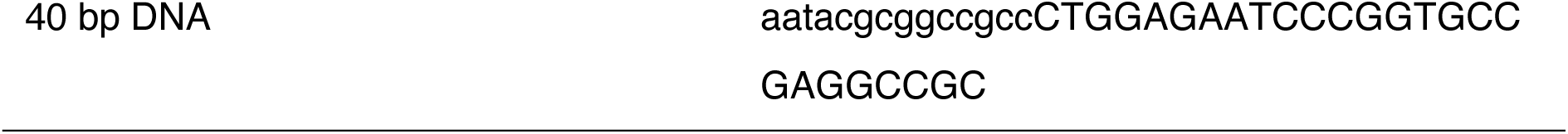
Constructs used in this study.

