## Supplementary figures and images for "Disease mutations in the PWWP domain of DNMT3A affect chromatin recruitment through multiple mechanisms"

### Supplemental Figure 1

##### Supp Figure 1

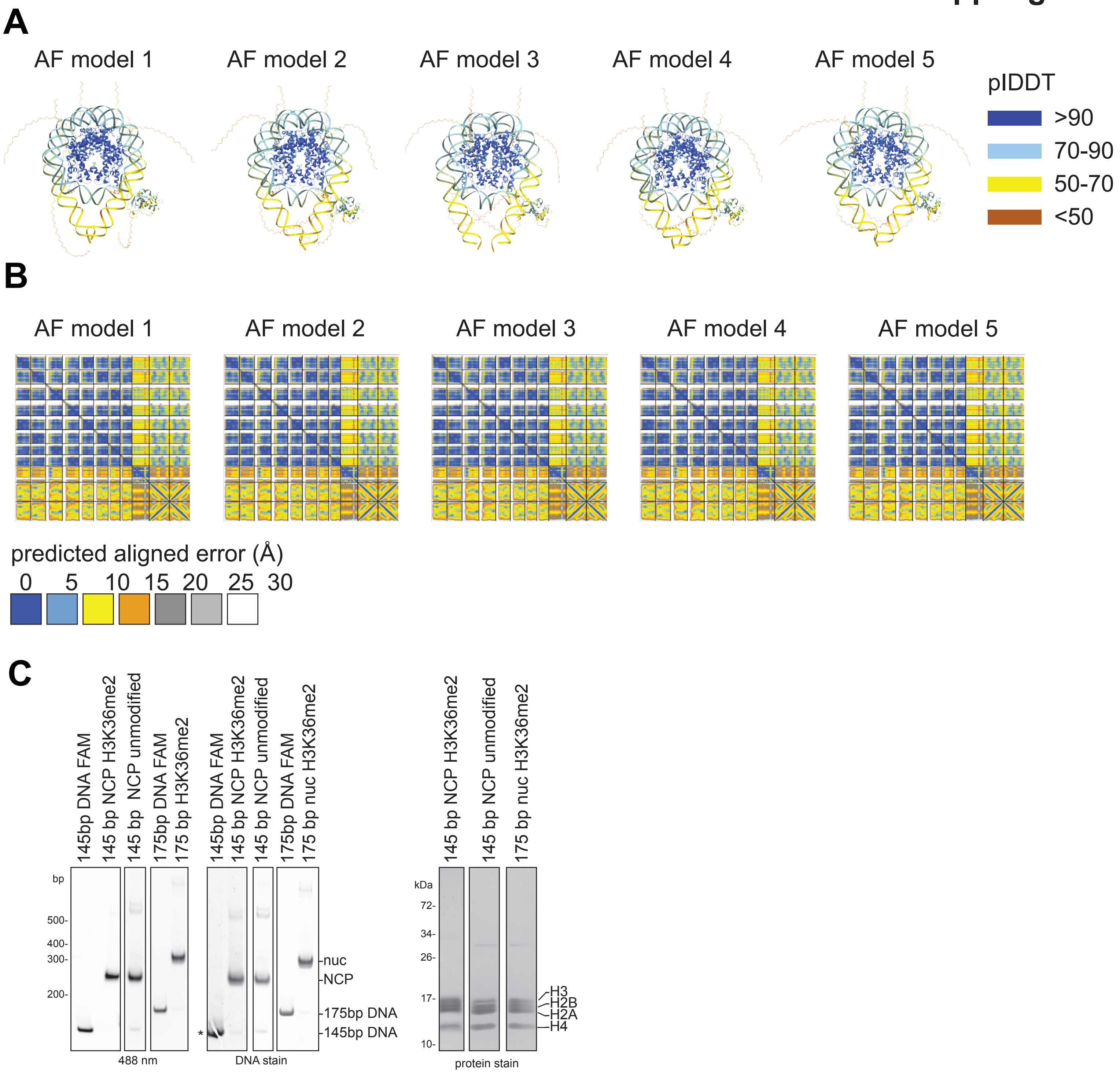

### Supplemental Figure 2

#### Supp Figure 2

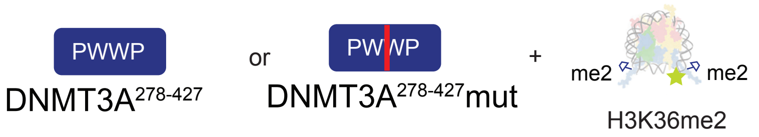


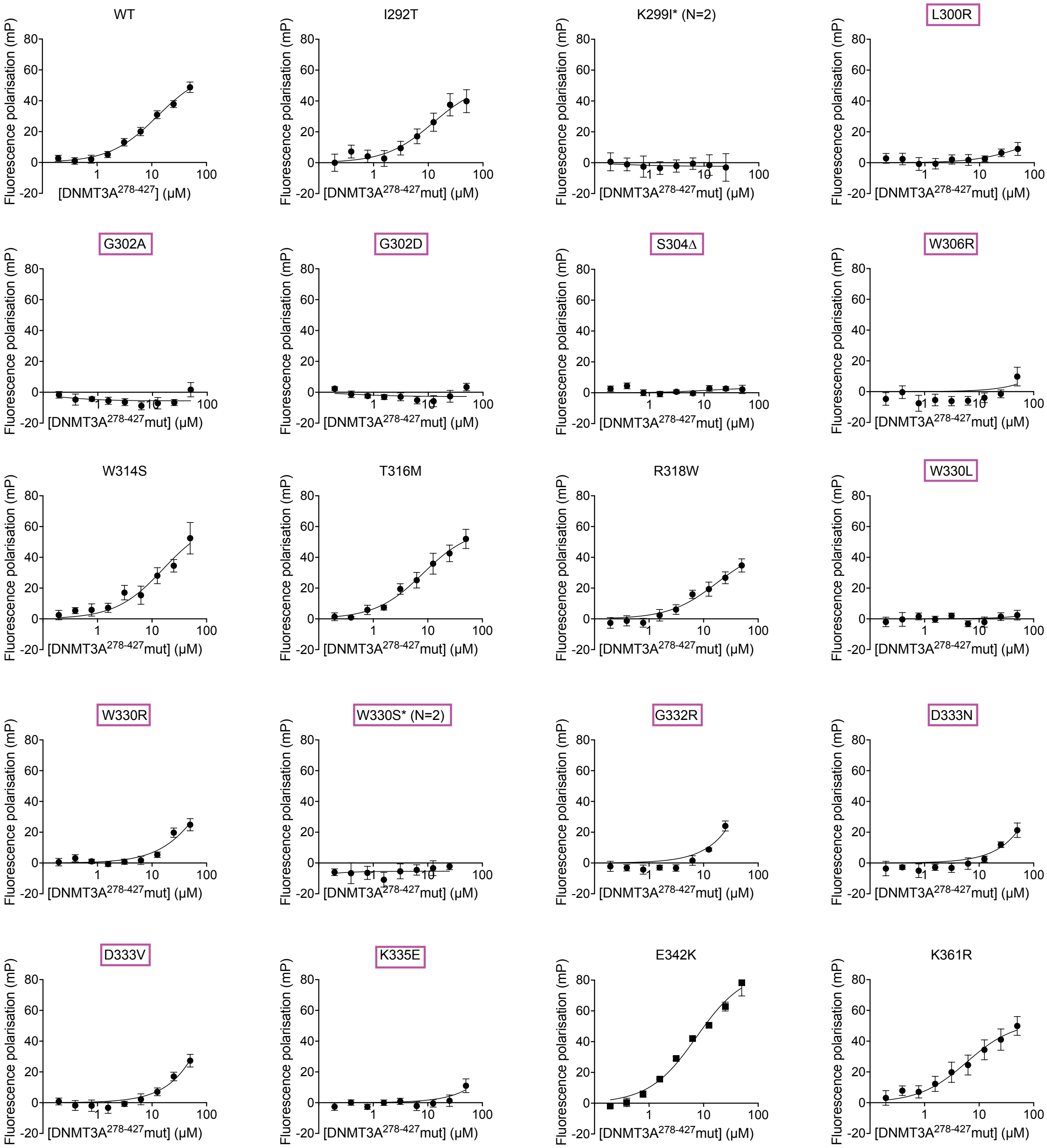

### Supplemental Figure 3

#### Supp Figure 3

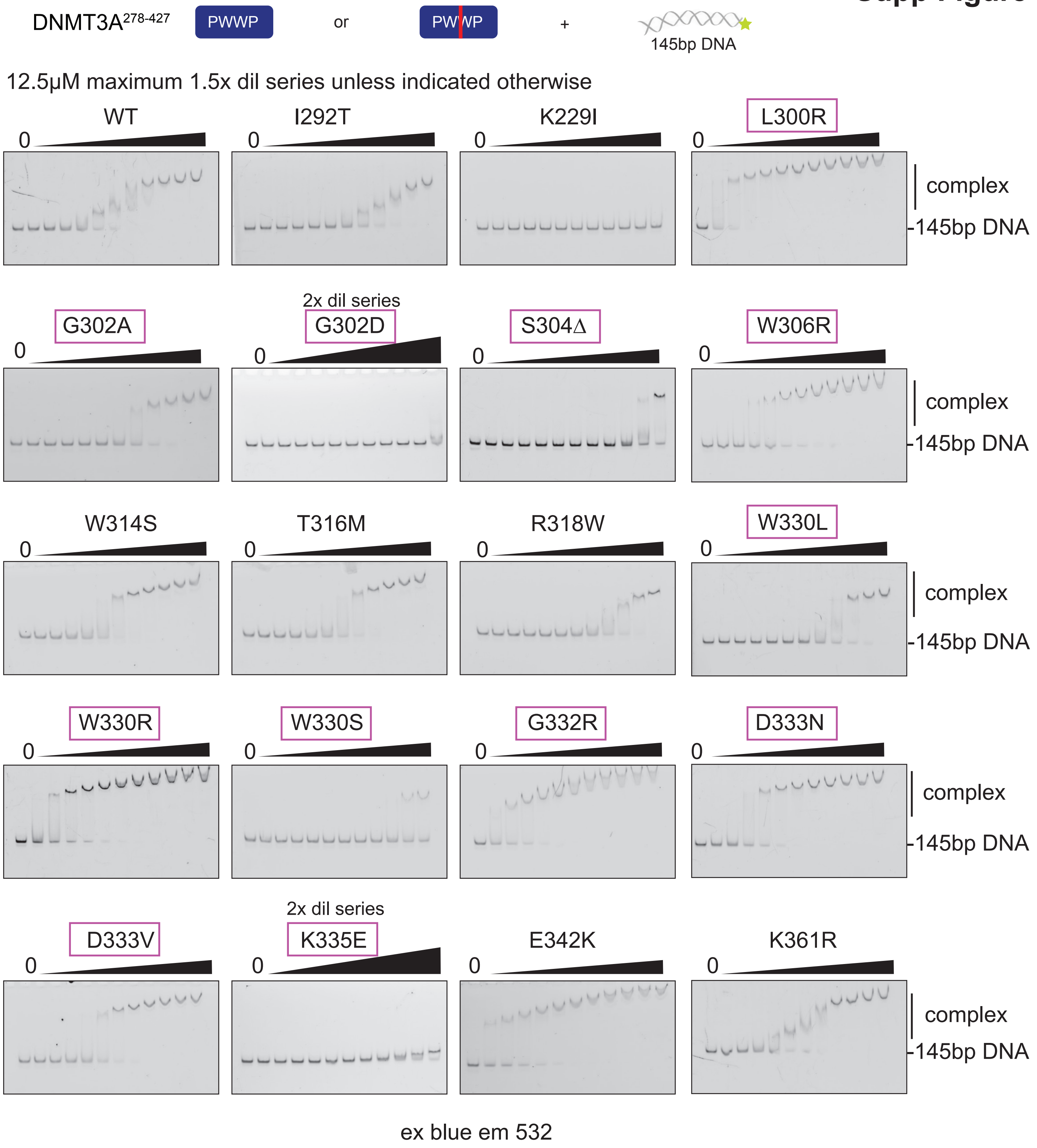

### Supplemental Figure 4

#### Supp Figure 4

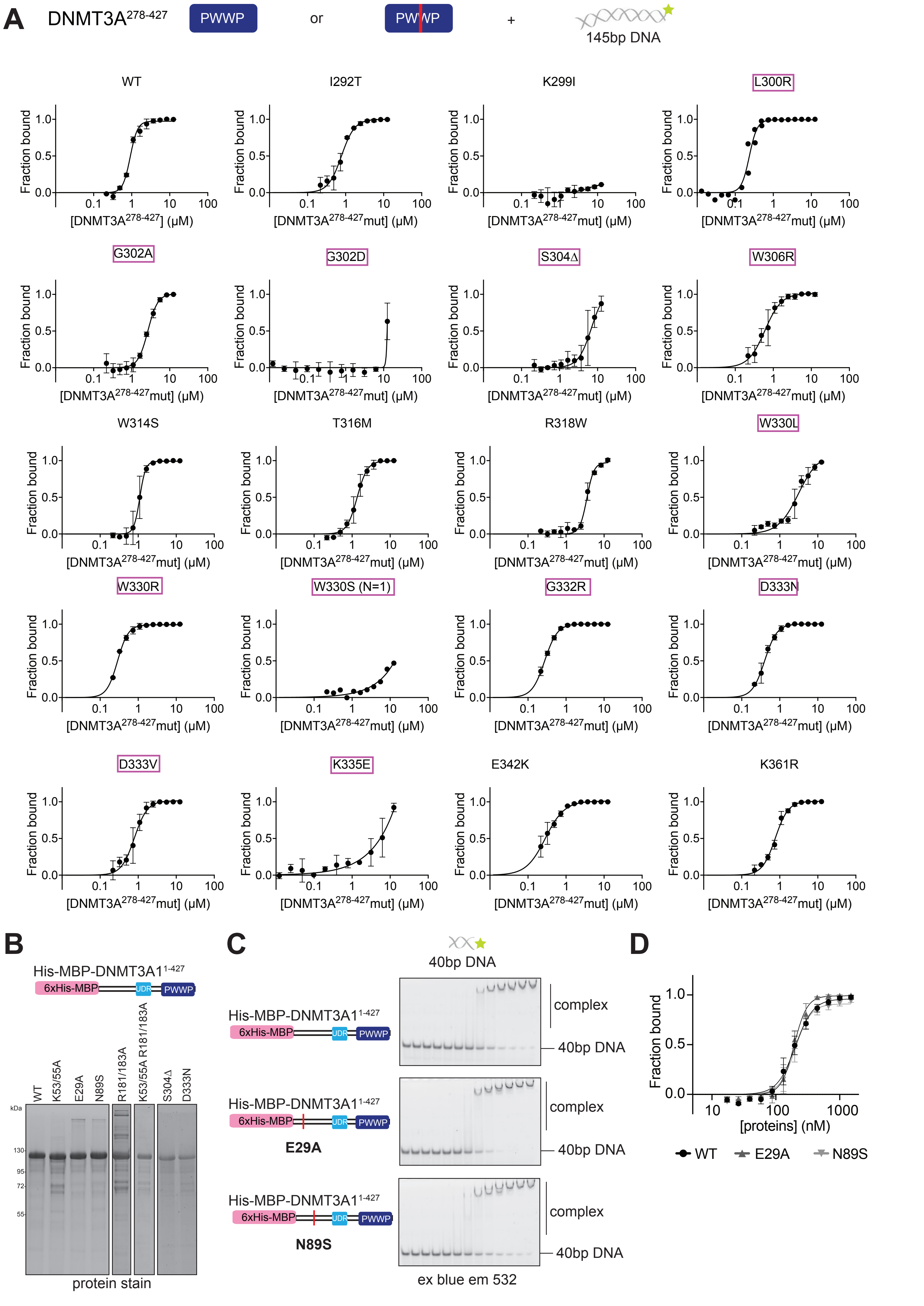

### Supplemental Figure 5

#### Supp Figure 5

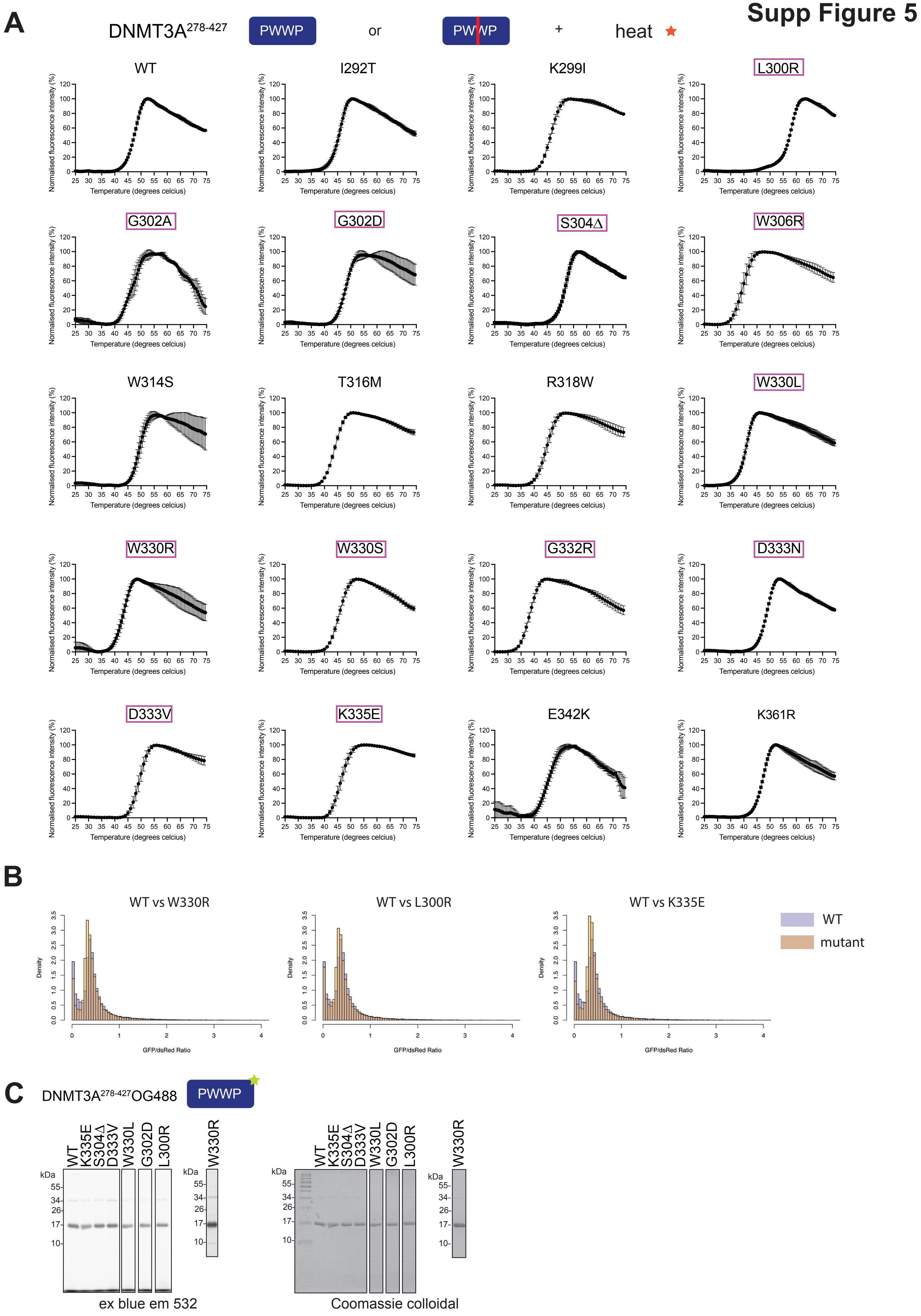
